# Senataxin modulates resistance to cisplatin through an R-loop mediated mechanism in HPV-associated Head and Neck Squamous Cell Carcinoma

**DOI:** 10.1101/2024.02.22.581374

**Authors:** Hannah Crane, Ian Carr, Keith D Hunter, Sherif F. El-Khamisy

## Abstract

**Introduction:** Oropharyngeal Squamous Cell Carcinoma (OPSCC) is a site defined subtype of head and neck cancer with two distinct clinical subtypes: HPV-associated (HPV+) and HPV-independent (HPV-); both of which are commonly treated with chemoradiotherapy involving cisplatin. Cisplatin creates DNA crosslinks, which lead to eventual cell death via apoptosis. Clinical outcomes in HPV-OPSCC are poor and although HPV+ has an improved response to therapy, a subset of patients suffer from distant metastases, with a poor prognosis. Therefore, there is a need to understand the molecular basis underlying treatment resistance. A common mechanism of chemotherapy resistance is upregulation of DNA repair, and a major source of endogenous DNA damage are DNA/RNA hybrids, known as R-loops. R-loops are three stranded DNA/RNA hybrids formed in the genome as a by- product of transcription and are normally transient; however, they can persist and become a source of genomic instability. The contribution of R-loops to the development of cisplatin resistance in OPSCC is unknown.

**Methods:** HPV+ and HPV- cisplatin resistant cell lines were developed, and RNA-sequencing was used to investigate changes in gene expression. Changes in R-loop dynamics were explored using slot blots and DRIP-qPCR. The effect of depleting known R-loop regulators on cisplatin sensitivity was assessed using siRNA. R-loop burden in a cohort of HPV+ and HPV- OPSCC tumours was explored using S9.6 immunohistochemistry.

**Results:** Development of cisplatin resistant clones led to changes in gene expression consistent with resistance, alongside alterations in the expression of known R-loop regulators. Both HPV+ and HPV- resistant cells had elevated global R-loop levels and in HPV+ resistant cells there was a corresponding upregulation of the R-loop resolving protein, senataxin, which was not observed in HPV- resistant cells. Depletion of senataxin led to increased sensitivity to cisplatin in both HPV+ and HPV- resistant cells, however, the effect was greater in HPV+ cells. Quantification of R-loop levels by S9.6 immunohistochemistry revealed that HPV+ tumours and tumours with bone metastases had a higher R-loop burden.

**Conclusion:** R-loops are involved in modulating sensitivity to cisplatin and may represent a potential therapeutic target.

## Introduction

Oropharyngeal Squamous Cell Carcinoma (OPSCC) is a site defined subset of head and neck cancer occurring in the base of tongue, soft palate and tonsils ^1^ which has shown a dramatic increase in incidence in the western world ^2–5^. Over the past three decades, the role of high risk Human Papilloma Virus (HPV), in a subset of OPSCC has become apparent ^6,7^. Given the overwhelming evidence of a causative role of HPV in OPSCC, in 2017 the World Health Organisation (WHO) categorised OPSCC into two distinct clinical subtypes, HPV-positive (HPV-associated/HPV+) and HPV-negative (HPV-independent/HPV-) ^1,8^. This categorisation was undertaken due to the markedly improved prognosis of HPV+ OPSCC (3-year survival of 82.4%) when compared to HPV-OPSCC (3-year survival of 57.1%) ^6^.

Platinum-based chemotherapeutic agents such as cisplatin are a mainstay in the treatment of OPSCC, alongside surgery and radiotherapy. Cisplatin (cis-diamminedichloroplatinum(II)) was first identified as an anti-tumour agent in 1969 ^9^ and following cisplatin uptake into a tumour cell, the low chloride concentration within the cytoplasm allows for replacement of chloride ions with water ^10^. This “aquated” cisplatin is able to bind to nuclear and mitochondrial DNA with high affinity, causing intra- and inter-strand DNA crosslinks, blocking transcription pathways and leading to eventual cell death by apoptosis ^11,12^.

OPSCC is commonly treated with chemoradiotherapy involving cisplatin or surgery followed by post-operative chemotherapy if there are adverse features present on histopathological examination of the surgical specimen ^13^. A systematic review has shown that chemotherapy, when used alongside surgery and radiotherapy, is associated with improved survival in patients with oral and oropharyngeal cancer ^14^. Furthermore, in HPV+ OPSCC, a trial of treatment de-escalation through replacement of cisplatin with the monoclonal antibody cetuximab resulted in inferior clinical outcomes ^15^. Therefore, platinum based chemotherapies, such as cisplatin, continue to be an important component of treatment for patients with HPV+ and HPV- OPSCC, improving clinical outcomes.

Notwithstanding the importance of cisplatin in the treatment of OPSCC, resistance to treatment is common, especially in HPV- OPSCC which persists with poor overall survival ^6^. Despite the overall improved prognosis of HPV+ OPSCC, there are a subset of patients who present with loco-regional recurrences or distant metastases, with a poor prognosis ^16–19^.

Additionally, a systematic review highlighted that patients with HPV+ OPSCC were more likely to undergo distant metastases to multiple organs when compared to HPV- OPSCC ^20^. Therefore, it is important to understand the molecular basis of cisplatin resistance in both HPV+ and HPV- OPSCC, in order to improve patient outcomes.

Resistance to cisplatin is multifactorial, and researchers have demonstrated the importance of a number of factors including tumour heterogeneity, altered cellular uptake and the effect of the surrounding tumour microenvironment ^21^. As cisplatin exerts it’s effects through formation of intra-and inter-strand crosslinks, the DNA damage response is also known to be crucial in mediating resistance to cisplatin therapies ^22^.

One of the major sources of endogenous DNA damage are DNA/RNA hybrids (R-loops). R-loops were first described in 1979 and exist as three stranded nucleic structures comprised of a DNA:RNA hybrid with an associated free strand of DNA ^23^. R-loops form as a by-product of transcription and next-generation sequencing studies have shown they occupy 5-10% of the genome ^24,25^, preferentially forming in regions of the genome which are G-rich or show GC-skew ^26,27^. R-loops are known to have a number of physiological roles, including enabling transcriptional activation through prevention of DNA methylation at promoter regions ^28^.

However, unscheduled or persistent R-loops are a potential source of genomic instability, largely due to their potential to cause transcription-replication conflicts ^29^. R-loops have been implicated in the pathogenesis of a number of tumours, including Embryonal Tumours with Multilayered Rosettes (ETMR) ^30^, Ewing Sarcoma ^31^ and Kaposi’s sarcoma ^32^. However, the contribution of R-loops to the development of cisplatin resistance in OPSCC has not previously been explored in the literature.

We hypothesised that R-loop physiology would change upon the development of cisplatin resistance and that it could be modulated for therapeutic benefit. To investigate this, we developed cisplatin resistant clones of a HPV+ and HPV- cell line and utilised these to explore R-loop dynamics upon the development of resistance.

## Materials and Methods

### Cell culture

Monolayer cell culture was undertaken in a class two biological cabinet and cells were maintained in 37°C incubators with 5% CO_2_. Two head and neck squamous cell carcinoma cell lines were used in this study, UPCISCC89 (HPV- ) and UDSCC2 (HPV+), henceforth referred to as SCC89 and SCC2. The presence of high risk HPV16 in the SCC2 cell line was confirmed by testing on a Roche Cobas 6800 instrument and through qPCR for E6 and E7 (Supplementary figure 1A-D). Both cell lines were STR profiled by NorthGene (Newcastle, UK) and regular mycoplasma testing was undertaken by the core facility service at the School of Clinical Dentistry, University of Sheffield. Cells were routinely cultured in low-glucose Dulbecco’s Modified Eagle’s Medium (DMEM), supplemented with 10% fetal bovine serum (FBS), 1% penicillin-streptomycin and 1% L-glutamine.

### Generation of cisplatin resistant clones

Cisplatin resistant clones were developed using methods previously described ^33–35^. Cisplatin was purchased from Sigma-Aldrich (Cis-Diammineplatinum(II) dichloride, P4394) and was dissolved in 0.9% Sodium Chloride (NaCl) to create a stock concentration of 1mM. This was aliquoted and stored at -20°C. SCC89 and SCC2 were treated long-term with cisplatin over a period of 2-3 months. Occasional cisplatin free passages were undertaken to allow cells to recover. SCC89 cells were grown in standard media supplemented with 2.5□M cisplatin, which increased to 5□M cisplatin after 2 passages. SCC2 cells were grown in standard media supplemented with 5□M cisplatin, which increased to 10□M cisplatin after 2 and 4 passages. Single cell clones were selected using serial dilution in a 96 well plate format and a single high dose of cisplatin (5□M for SCC89 and 10□M for SCC2).

### Antibodies, siRNA and primers

Details regarding the antibodies, siRNA and primers used in this study can be found in supplementary tables 1-3. siRNA transfections were undertaken using Lipofectamine RNAiMAX, following the manufacturer’s instructions. USP11 siRNA was used at a final concentration of 20nM and pooled senataxin siRNA was used at a final concentration of 80nM.

### RNA-Sequencing

RNA was extracted using the Monarch Total RNA miniprep kit (New England Biotechnologies (NEB) #T2010) or the Qiagen RNeasy mini kit (74104) according to the manufacturer’s instructions. RNA was quantified using Nanodrop 100 Spectrophotometer (ThermoFisher Scientific) and quality assured using A_260:280_ ratio. cDNA was prepared using the Applied Biosystems High-Capacity cDNA Reverse transcription kit according to the manufacturer’s instructions.

RNA from the parental cells and subsequent resistant clones was sent to Novogene (Cambridge, UK) for mRNA sequencing (polyA library prep, 20 million paired reads per sample, 150bp paired-ended sequencing) on an Illumina NovaSeq. The resulting fastq files were quality assessed using fastQC and combined using multiQC ^36^. Following quality assessment, the fastq files were aligned and quantified using Salmon ^37^ on the high-performance computer (HPC) cluster at the University of Sheffield, using Genome Reference Consortium Human Build 38 (GRCh38). The resulting transcript quantification files were imported into the statistical programme R (version 4.2.3) and differential expression was conducted using DESeq2 ^38^. The cut-off for differential expression was set as an adjusted p-value less than 0.05 with a log2fold change of greater than 1. Resulting differentially expressed genes underwent gene ontology analysis using clusterProlifer ^39^.

### Quantitative PCR (RT-qPCR)

cDNA was diluted 1:5-1:20 with nuclease-free water, dependent on the input amount of RNA. Dilutions were kept consistent within the same experiment. Each qPCR reaction contained 5□l of diluted cDNA, 2.8□l of forward and reverse primer at 5□M concentration and 10□l QuantiNova (Qiagen), made up to a final volume of 20□l with nuclease-free water. Serially diluted standards were run with each reaction. Reactions were run on a Rotor-Gene 6000 qPCR machine (Qiagen) and the following thermocycling conditions were used: denaturation at 95°C for 10 minutes, followed by 50 cycles of denaturation at 95°C for 15 seconds, annealing at 5°C below the average melting temperature of the forward and reverse primer for 15 seconds and extension at 72°C for 30 seconds. A melt curve was added to the end of each reaction by increasing the temperature from 72°C to 95°C in 1°C increments at 5 second intervals. CT values were determined for each reaction using the Q-Rex software and analysed using the delta-delta CT method, normalising gene expression to Actin or GAPDH for each reaction.

### Cell viability assays

MTS assays were undertaken using the CellTiter 96 Aqueous One Solution Cell Proliferation Assay (Promega) or Abcam MTS assay kit (ab197010). Briefly, cells were seeded at an appropriate density in 96 well plates and allowed to attach overnight. Following transfection and 24 hours of cisplatin treatment, an appropriate amount of MTS reagent was added to each well following the manufacturer’s instructions and the plate incubated at 37°C for 1-4 hours. The plate was subsequently read on using a Tecan spectrophotometer at 490nm.

Clonogenic assays were performed as previously described ^40^. Cells were seeded at a density of 3000 to 4000 cells on a 10cm dish and incubated overnight. The following day, cells were treated with the indicated concentrations of cisplatin and left to form colonies for 7-10 days. Following visible colony formation, the media was removed, plates washed with PBS and subsequently fixed with 80% ethanol for 15 minutes. The plates were then air dried for 5 minutes and then stained with 0.5% crystal violet for 30 minutes. The number of colonies on each plate was counted using an automated cell counter (Protos 3) and the surviving fraction was calculated by dividing the number of colonies on treated plates by the number of colonies on untreated plates.

### Western blotting

Cells were harvested and protein was extracted using lysis buffer (20mM HEPES pH7.4, 2mM MgCl_2_, 40mM NaCl and 1% Triton-X) containing complete mini EDTA protease inhibitor (Roche) at 1 in 50 concentration, BaseMuncher (Abcam) at 1 in 1000 concentration and where appropriate PhosSTOP (Sigma) at a final concentration of 1 in 20. 30-50□l of lysis buffer was added to the cell pellet and left on ice for 20 minutes with regular vortexing. The samples were centrifuged at 13000rpm for 15 minutes and the resulting supernatant was stored at -20°C. The protein was quantified with the Bradford Technique, using Coomassie Blue (Thermofisher). Equal amounts of protein were diluted in 5x protein loading buffer and boiled at 95°C for 5 minutes before being loaded onto an SDS-PAGE gel. Gels were either pre-cast 4-15% gels (Invitrogen) or a 4-20% gradient gel was prepared in glass cassettes ^41^.

Samples were run at 120V for 10 minutes, followed by 170V for 1 hour. Gels were transferred onto nitrocellulose membrane using a Trans-Blot Turbo (Bio-Rad), using the pre-set high molecular weight setting. Following transfer, membranes were blocked for 1 hour at room temperature with 5% dried milk in Tris-buffered saline with 0.1% Tween (TBST) and subsequently incubated with primary antibody overnight at 4°C. The following primary antibodies were used: Senataxin (1:1000 dilution, Bethyl, A301 – 104A or 105A), USP11 (1:1000 dilution, Bethyl, A301 – 613A), and Beta-Actin (1:1000, Sigma, A5316). Following three 5-minute TBST washes, blots were incubated with an appropriate mouse or rabbit HRP secondary antibody (Bio-Rad) at 1:4000 concentration for 1 hour at room temperature. Following three further TBST washes, blots were visualised using Clarity Western ECL substrate (Bio-Rad), on a ChemiDoc Imaging system (Bio-Rad) and quantified using Image Studio Lite (Licor).

### Slot blotting

DNA was extracted using a phenol-chloroform method (described below) or with a DNeasy Blood and Tissue Kit (Qiagen), following the manufacturer’s instructions. 400ng or 5μg of DNA was dotted onto a nylon or nitrocellulose membrane respectively using a using a slot blot apparatus (Hoefer), as per the manufacturer’s instructions. Where indicated, samples were pre-treated with exogenous RNase H (M0297, NEB), following the manufacturer’s instructions. Following the samples being applied to the membrane, if a ssDNA antibody was used then the blot was denatured for 10min in 1.5M NaCl, 0.5M NaOH solution and then neutralised for 10min in 0.5M Tris-HCl pH 7.0, 1M NaOH solution. This step was omitted for use of the dsDNA antibody. The blots were UV crosslinked (120000uJ/cm^2^), blocked in 5% dried milk powder in TBST for 1 hour at room temperature and subsequently incubated with primary antibody overnight at 4°C. The following primary antibodies were used: S9.6 (1:500 dilution, isolated from hybridoma by BioServ (Sheffield, UK)), ssDNA (1:5000 dilution, EMD Millipore, MAB3868) and dsDNA (1:500 dilution, Santa-Cruz, sc-58749). Following incubation with the primary antibody, blots were washed three times in TBST for 5 minutes each and incubated with an anti-mouse HRP secondary (Bio-Rad) at 1:2000 dilution for 1 hour at room temperature. Following three final 5-minute washes with TBST, blots were visualised and quantified as detailed in the western blot section above.

### DNA:RNA immunoprecipitation (DRIP-qPCR)

DRIP was undertaken based on method previously published by Sanz and Chédin ^42^. Briefly, DNA was extracted as follows. Cells were harvested and resuspended in 1.6ml TE buffer, pH8 (Thermofisher) with 50□l of 20% SDS and 10□l of 10mg/ml of proteinase K and incubated overnight at 37°C. The lysate was added to a MaXtract high density tube with an equal amount of phenol/chloroform/isoamyl alcohol and mixed by inversion 5 times.

Following centrifugation at 1500g for 10 minutes, the clear supernatant was added to a 15ml tube containing 4ml of 100% ethanol and 160□l of 3M sodium acetate. The DNA was precipitated by mixing on a rotary mixer at 10rpm for 10 minutes and the DNA was spooled out and washed with 80% ethanol three times for 10 minutes each. The DNA was then air dried and resuspended in 125□l of TE buffer. A restriction enzyme digest was then set up to incubate overnight at 37°C containing up to 118.5□l of DNA, 1.5□l of 10m/ml BSA (NEB), 15□l of 2.1 buffer (NEB), 30 units of SSP1 (NEB), 30 units of BSRG1 (NEB), 30 units of ECoR1 (NEB), 30 units of HindIII (NEB), 30 units of XBA1 (NEB), 1.5□l of 100mM spermidine and the reaction was made up to 150□l with nuclease free water. The following day the DNA was added to a phase lock light gel tube (Quantabio), with 100□l of nuclease free water and 250□l of Ultrapure^TM^ phenol:chloroform:isoamyl alcohol (Thermofisher). Following centrifugation at 16000g for 10 minutes, the supernatant was added to 625□l 100% ethanol, 25□l sodium acetate and 1.5□l of glycogen and incubated at -20°C for at least one hour. This was followed by a centrifugation at 16000g for 35 minutes at 4°C. The supernatant was removed and 200□l of 80% ethanol was added followed by a further centrifugation at 16000g for 10 minutes at 4°C. The supernatant was removed, the DNA pellet was air-dried, resuspended in 50□l of TE buffer, quantified using a Nandrop 100 spectrophotometer and stored at -80°C.

For each condition, 8μg of DNA was diluted in 500□l of TE buffer and 50□l was taken as input. For the RNase H treated sample, an equal amount of DNA (8μg) was treated with 8□l of RNase H in 1x RNase H buffer (NEB) for 5 hours at 37°C. Following RNase H digestion, 400□l of TE buffer was added to the RNase H sample and subsequently treated identically to the non-RNase H treated sample. 52□l of 10x DRIP binding buffer (2.8ml 5M NaCl, 1ml 1M sodium phosphate and 50□l Triton-X in 10ml nuclease free water) and 10□l of S9.6 antibody were added to each sample (RNase H treated and untreated) and incubated for 14-17 hours with 10rpm rotation at 4°C. The following day, 90□l of Protein G beads were washed twice with 700□l 1x DRIP binding buffer for 10 minutes at 10rpm. The DNA was added to the washed beads and incubated for 2 hours at 4°C with 10rpm rotation. The supernatant was discarded, and the beads were washed twice with 750□l of 1X DRIP binding buffer for 15 minutes with 10rpm rotation. 300□l of DRIP elution buffer was then added to the beads with 14□l of 10mg/ml proteinase K and incubated at 55°C for 45 minutes with 10rpm rotation. The resulting supernatant was transferred to a phase lock light gel tube (Quantabio) and phenol-chloroform purification undertaken as detailed in the previous paragraph. The resulting DRIP-DNA was stored at -80°C until qPCR.

DRIP-qPCR was undertaken using QuantiNova SYBR Green-based PCR (Qiagen) on a Rotor-Gene 6000 (Qiagen). Standards were produced by pooling equal amounts of each input and serially diluting 1:10. Each reaction contained 10□l of QuantiNova, 2.8□l of forward and reverse primer pair (at 5□M concentration), 2□l of DNA and 5.2□l of nuclease free water.

The cycling conditions used were the same as those used for RT-qPCR and the primers used are detailed in Supplementary Table 1.

### Immunofluorescence

Cells were seeded at an appropriate density onto 13mm glass coverslips in a 24 well plate. Following transfection and cisplatin treatment for the time indicated in each figure, the media was removed, and wells washed with 500□l of PBS. Cells were then fixed with 3.7% formaldehyde for 10 minutes at room temperature, followed by permeabilisation with 0.5% Triton-X for 5 minutes. Wells were then washed with 500□l of PBS three times, followed by blocking with 500□l of 3% BSA in PBS-T for 1 hour at room temperature. Wells were incubated with 160□l of primary antibody diluted in 3% BSA for 1 hour at room temperature or overnight at 4°C as follows: Senataxin (1:1000 dilution, Bethyl, A301 – 104A or 105A), USP11 (1:500 dilution, Bethyl, A301 – 613A), γH2AX (1:1000, EMD Millipore 05 – 636).

Following three washes with 500□l of PBS-T the wells were incubated with the following secondary antibodies in the dark for 1 hour at room temperature at 1:500 dilution: Alexa Fluor^TM^ 488 anti-mouse (a11001), Alexa Fluor^TM^ 488 anti-rabbit (a11008) and Alexa Fluor^TM^ 555 anti-rabbit (a21428). Following two washes with 500□l PBS-T and one wash with 500□l PBS the coverslips were removed and allowed to dry for 10 minutes at room temperature before mounting with Immun-Mount^TM^. Visualisation was undertaken on Zeiss Axioplan 2 microscope and images were taken using a 100x objective. The resulting images were analysed in ImageJ using a pre-defined macro to determine the mean nuclear intensity for each cell.

### Alkaline single cell gel electrophoresis (comet) assay

Following treatment, cells were harvested and resuspended at 300,000 cells/ml in PBS and mixed with an equal amount of 1.2% low melting point agarose. This was placed underneath a coverslip on top of pre-prepared 0.6% agarose and allowed to set at 4°C for 1 hour. Once set, the coverslips were removed and 1ml of lysis buffer was added to each slide (2.5M NaCl, 10mM Tris HCl, 100mM EDTA pH8.0, 1% Triton X-100, 1% DMSO; pH10) and incubated for 1 hour in the dark at 4°C. Alkaline electrophoresis buffer was prepared by adding 2ml 0.5M EDTA, 5ml 10M NaOH and 10ml DMSO with dH_2_0 to a final volume of 1L. Following lysis, slides were placed in the comet tank with the alkaline electrophoresis buffer for 45 minutes. The comet tank was then run at 12V for 25 minutes and slides were removed and neutralised with 1ml of 0.4M Tris pH7 overnight at 4°C. The slides were visualised on a fluorescence microscope with 1:10,000 SYBR green (Sigma) and quantified using Comet Assay IV. At least 100 cells were scored for each biological replicate and the mean comet tail was analysed.

### Immunohistochemistry

Ethical approval was sought and obtained to undertake immunohistochemistry on human tissue samples (19/YH/0029). Following sectioning, tissue was mounted on SuperFrost Plus^TM^ slides (Epredia) and baked at 60°C for 60 minutes. Slides were dewaxed in xylene twice for 5 minutes each, followed by two 5 minutes incubation in 100% ethanol.

Endogenous peroxidase activity was blocked by incubation with 3% H_2_0_2_ in methanol for 20 minutes. Slides were briefly washed in PBS, followed by antigen retrieval using sodium citrate buffer in a steamer for 30 minutes (10mM Sodium citrate, 0.05% Tween-20, pH6.0). Where indicated, after antigen retrieval but prior to endogenous blocking, slides were treated with RNase H (25 units RNase H in 1x RNase H buffer) or mock treated with buffer only overnight at 37°C in a humidity chamber filled with dH_2_0. Slides were washed in PBS and a hydrophobic barrier was drawn around the slides with ImmEDGE^TM^ Hydrophobic Barrier Pen (Vector Laboratories). Slides were blocked in 100% horse serum for 1 hour at room temperature in a humidity chamber. The primary antibody was diluted in 100% horse serum (S9.6 at 1:1500 dilution) and slides were incubated overnight at 4°C in a humidity chamber. Following two washes in PBS for 5 minutes each, secondary antibody and ABC complex (VECTSTAIN® Elite® ABC-HRP Kit, Peroxidase (Mouse IgG)) were added according to the manufacturer’s instructions. Following two further washes in PBS for 5 minutes each, staining was visualised using DAB Substrate Kit, Peroxidase (HRP) (Vector Laboratories) for 7 minutes. The reaction was ceased with dH_2_0 and the slides counterstained with haematoxylin using a Lecia automated stainer. Slides were mounted using DPX mountant and visualised on an Olympus light microscope. Images were taken using cellSens image software and scale bars on each image indicate the magnification (200□M: 4x magnification, 100□M: 10x magnification, 50□M: 20x magnification, 20□M: 40x magnification). Where indicated, slides were scanned using a Leica Aperio CS2 and saved as *.svs files. Images and digital slides were analysed using QuPath (Version 0.3.0) ^43^.

### Statistical analysis

Statistical analysis was carried out in GraphPad Prism 9 or R (version 4.2.3 "Shortstop Beagle”) using statistical analysis as detailed in each figure legend. The data presented are the mean and the standard error of the mean (sem) of three biological replicates, unless otherwise stated.

## Results

### Long term treatment with cisplatin results in resistant clones which show differential expression of R-loop regulators

In order to investigate platinum resistance, cisplatin resistant cells were developed using long term cisplatin treatment of a HPV+ and HPV- cell line, which led to a statistically significant increase in the half maximal inhibitory concentration (IC50) in both cell lines (Figure 1A-B). Following selection of single cell clones, clonogenic assays were performed to confirm resistance and established that the selected clones were more resistant to cisplatin treatment compared to the parental cells (Figure 1C-F).

**Figure 1:**
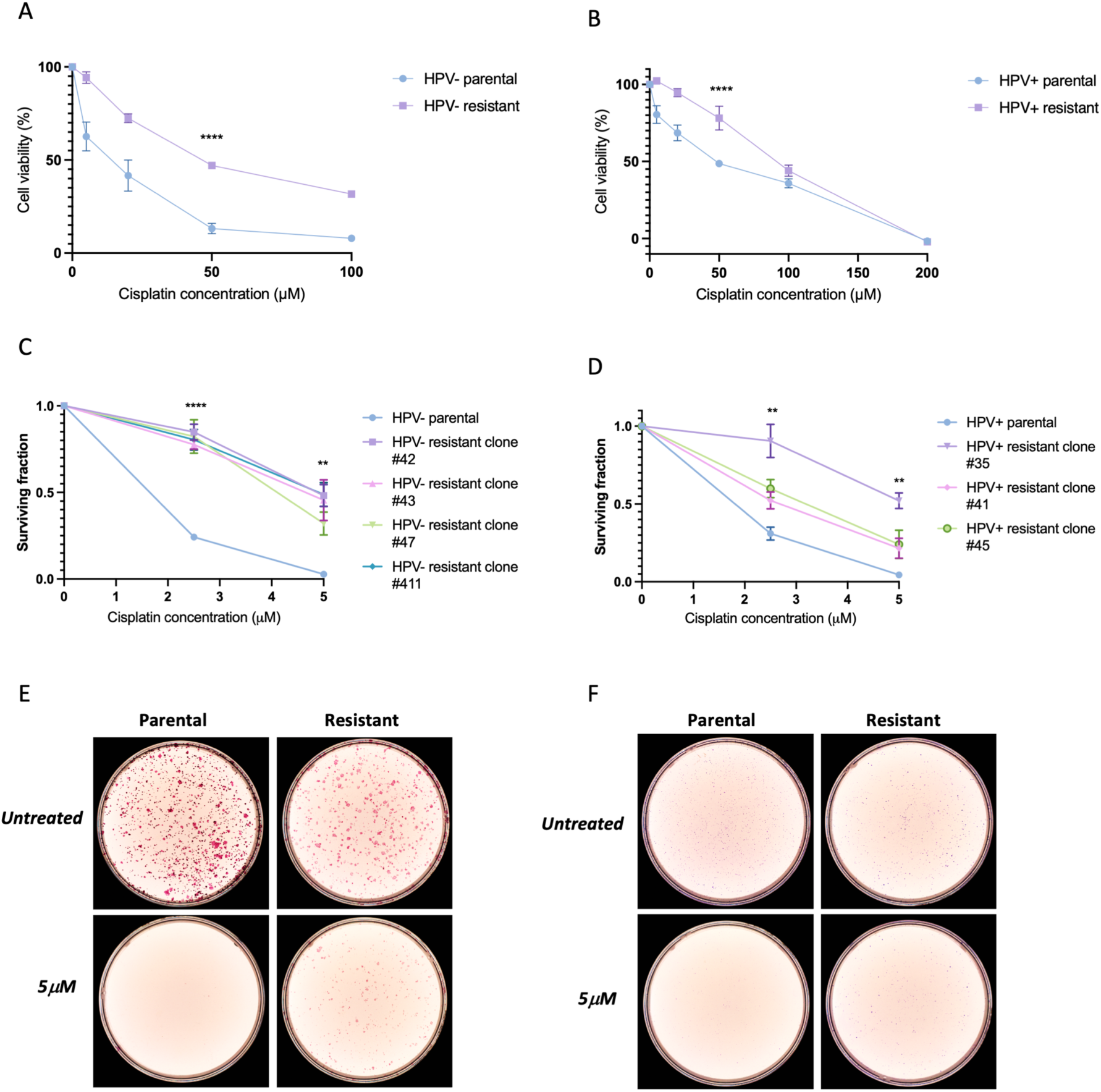
Long-term treatment with cisplatin results in the development of resistant clones. A. Cell viability assessed with an MTS assay following long-term treatment with cisplatin in HPV- (SCC89) parental cells (IC50: 10.0□M) and resistant cells (IC50: 47.6□M) (n=3, mean +/- sem). B. Cell viability assessed with an MTS assay following treatment with long term cisplatin in HPV+ (SCC2) parental cells (IC50: 40.0□M) and resistant cells (IC50: 84.9□M) (n=3, mean +/- sem). C. Results of clonogenic assay quantification to assess differences in surviving fraction of HPV- parental cells and resistant clones to cisplatin treatment. (n=3, mean +/- sem). D. Results of clonogenic assay quantification to assess differences in surviving fraction of HPV+ parental cells and resistant clones to cisplatin treatment. (n=3, mean +/- sem). E. Representative pictures of clonogenic assays with HPV- clone #411, treated with 5□M cisplatin. F. Representative pictures of clonogenic assays with HPV+ clone #35, treated with 5□M cisplatin. Statistical analysis carried out in A and B using non-linear regression and extra sum-of-squares F test was used to compare LogIC50 between datasets. Statistical analysis carried out in C and D using one way-ANOVA at both 2.5□M and 5□M concentrations with Tukey’s post-hoc test. ns=not significant, * = p<0.05, ** = p≤0.01, *** = p≤0.001, **** = p≤0.0001.

A selected resistant clone and associated parental cells were subjected to RNA-sequencing to explore transcriptome wide changes. Upon development of resistance, there were 1521 and 1234 differentially expressed transcripts in the HPV- and HPV+ cells respectively (Figure 2A-B). Two up-regulated and two down-regulated genes were validated for both the HPV+ and HPV- clones (Figure 2E-F), and successful validation was achieved for 7 of these targets. Differential expression of genes known to be involved in cisplatin resistance in other tumour types were identified, including up-regulation of Cyclic Nucleotide Gated Channel Subunit Beta 1 (CNGB1) which is associated with cisplatin resistance and reduced progression free survival in bladder cancer ^44^. Gene pathway analysis revealed a number of differentially expressed pathways (Figure 2C-D). The Wnt signalling pathway was the most over-represented pathway upon development of cisplatin resistance in HPV+ cells (Figure 2D), and this pathway has previously been associated with cisplatin chemoresistance in lung adenocarcinoma ^45^. Interestingly, the oxidative phosphorylation pathway was over-represented in both HPV+ and HPV- resistant cells (Figure 2C-D). Cisplatin treatment is known to increase levels of metabolites associated with generalised oxidative stress ^46–48^, and treatment with reactive oxygen species (ROS) scavengers increases resistance to cisplatin ^46,49^. Given that oxidative phosphorylation pathways were differentially expressed in resistant cells and ROS have previously been shown to increase R-loop levels ^50–52^, we next explored if there was differential expression of known R-loop regulators, using the publicly available database R-loopBase ^53^. R-loopBase is a database which utilises multiomics analysis and literature searching to collate tiers of known R-loop regulators, with the highest tier of regulator (tier 1) having been validated in multiple in-vitro assays ^53^. Using this database there were 134 and 62 differentially expressed R-loop regulators in the HPV- and HPV+ resistant cells respectively (Figure 2G). Furthermore, there were 5 and 3 differentially expressed tier 1 regulators in the HPV- and HPV+ cells respectively, with downregulation of RNASEH2C identified in both groups.

**Figure 2:**
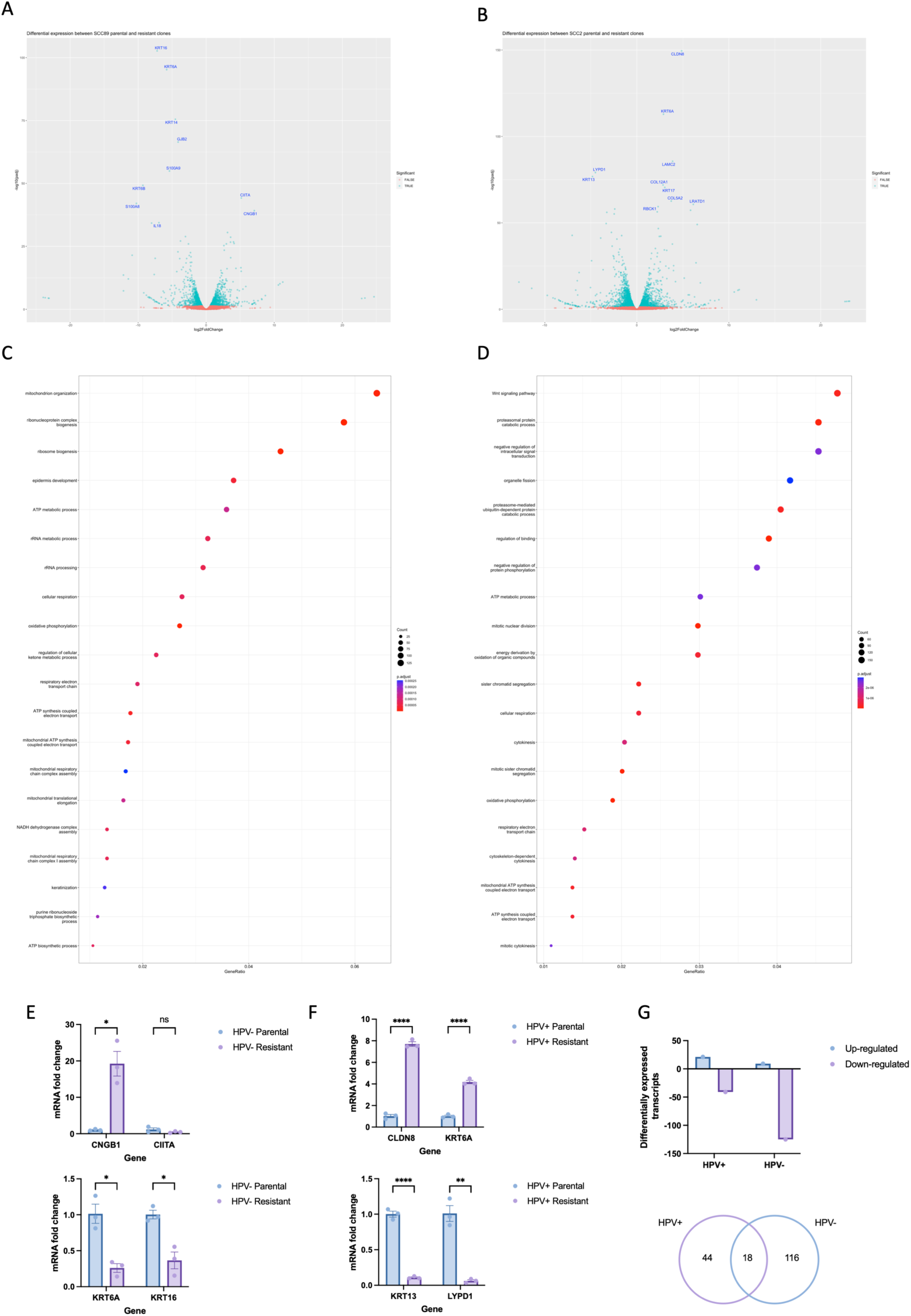
RNA-sequencing of the parental cells and associated resistant clone reveals differential expression of genes and pathways known to be involved in cisplatin resistance. A. Volcano plot demonstrating differentially expressed genes in the HPV- resistant clone compared to parental cells, with the top 10 differentially expressed genes labelled. B. Volcano plot demonstrating differentially expressed genes in the HPV+ resistant clone compared to parental cells, with the top 10 differentially expressed genes labelled. C. Gene expression analysis of differentially expressed genes in HPV- resistant cells compared to parental cells using clusterProlifer. D. Gene expression analysis of differentially expressed genes in HPV- resistant cells compared to parental cells using clusterProlifer. E. Validation of two up-regulated and two down-regulated transcripts in the HPV- resistant cells using RT-qPCR (n=3, mean +/- sem). F. Validation of two up-regulated and two down-regulated transcripts in the HPV+ resistant cells using RT-qPCR (n=3, mean +/- sem). G. Number of differentially expressed R-loop regulators in the HPV+ and HPV- resistant cells, with the overlap in differentially expression demonstrated with a Venn diagram. Statistical analysis conducted in E and F using multiple unpaired t-tests. ns=not significant, * = p<0.05, ** = p≤0.01, *** = p≤0.001, **** = p≤0.0001.

### Upon development of cisplatin resistance, HPV+ cells have an increase in global R- loops, with an associated upregulation of senataxin expression

The identification of differential expression of R-loop regulators upon development of cisplatin resistance led us to next explore global R-loop levels using an S9.6 slot blot. Initially the effect of cisplatin treatment on global R-loop levels in the cisplatin sensitive parental cells was investigated and we identified that treatment with 24 hours of cisplatin led to an increase in global R-loop levels in both HPV+ and HPV- cells (Figure 3A-D). A comparable increase in global R-loop level was identified following cisplatin treatment in HPV+ resistant cells, however in HPV- resistant cells there was no observed increase following cisplatin treatment (Figure 3A-D).

**Figure 3:**
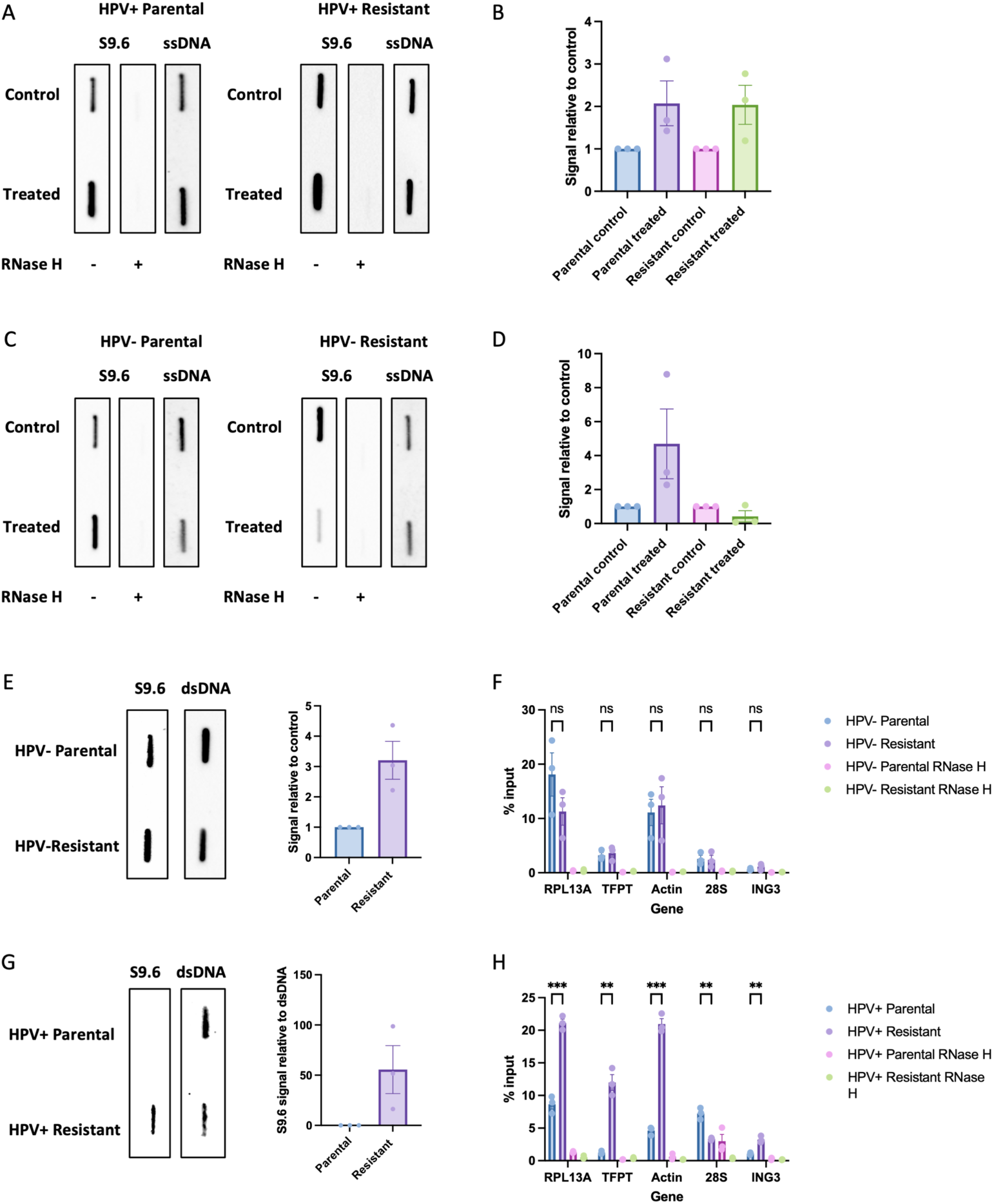
Upon development of cisplatin resistance, HPV+ and HPV- cells show alterations in R-loop dynamics. A. S9.6 slot blot showing global R-loop levels following treatment of HPV+ parental and resistant cells with 5□M cisplatin or vehicle only control for 24 hours. B. Quantification of three repeats of A (n=3, mean +/- sem). C. S9.6 slot blot showing global R-loop levels following treatment of HPV- parental and resistant cells with 5□M cisplatin or vehicle only control for 24 hours. D. Quantification of three repeats of C (n=3, mean +/- sem). In A and C, ssDNA (single-stranded DNA) was used as a loading control. E. S9.6 slot blot to evaluate global differences in R-loops at baseline between HPV- parental and resistant cells, with quantification of three repeats (mean +/- sem). F. Percentage input at positive R-loop loci in HPV- parental and resistant cells in untreated conditions, with associated RNase H treated controls (n=3, mean +/- sem). G. S9.6 slot blot to evaluate global differences in R-loops at baseline between HPV+ parental and resistant cells, with quantification of three repeats (mean +/- sem). dsDNA (double-stranded DNA) was used as a loading control in E and G. H. Percentage input at positive R-loop loci in HPV+ parental and resistant cells in untreated conditions, with associated RNase H treated controls (n=3, mean +/- sem). Statistics carried out in F and H using multiple unpaired t-tests (n=3, mean +/- sem). ns = not significant, ** = p≤0.01, *** = p≤0.001. For all figures, where indicated DNA was treated with RNase H sourced from NEB (M0297).

Investigation into changes in R-loop levels in untreated conditions revealed a global increase in R-loops in both HPV+ and HPV- cells upon the development of cisplatin resistance, with a larger increase noticed in the HPV+ cells (Figure 3E and 3G). However, when R-loop burden was investigated at specific genomic R-loop loci using DRIP-qPCR (*RPL13A, TFPT, Actin, and ING3*), we found an increase in R-loop occupancy in the HPV+ resistant cells (Figure 3H), which was not identified in the HPV- resistant cells (Figure 3F).

Senataxin is an helicase which is known to resolve R-loops at transcription termination sites ^54^. Interestingly, there was an upregulation of senataxin protein upon development of cisplatin resistance in HPV+ cells (Figure 4A-B), which was mirrored in an increase in senataxin mRNA levels (Figure 4C). Conversely, there was no increase in senataxin protein or mRNA levels in the HPV- resistant cells (Figure 4D-F).

**Figure 4:**
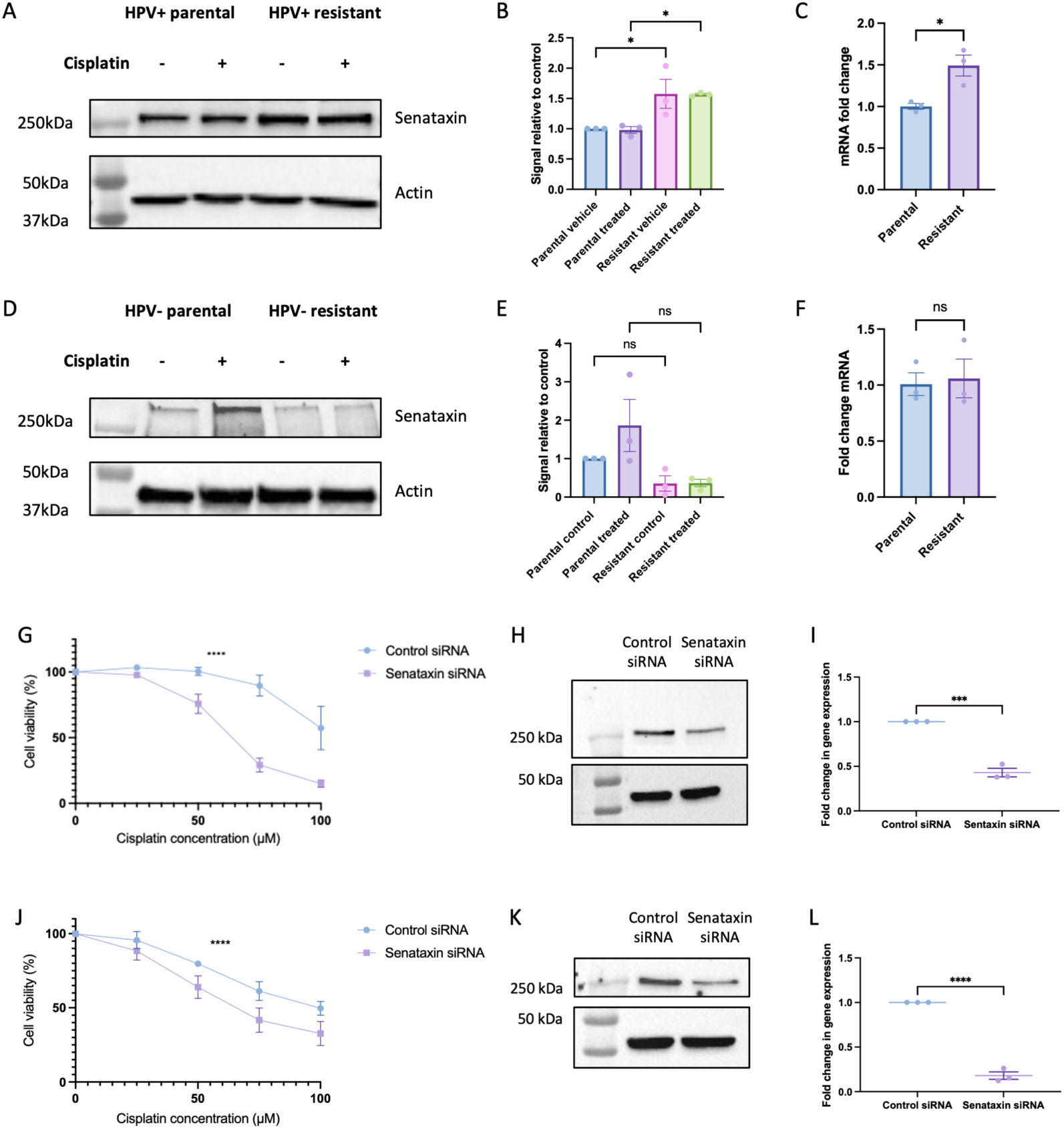
HPV+ resistant cells upregulate senataxin upon development of cisplatin resistance and showed reduced cell viability following cisplatin treatment in the presence of senataxin knockdown. A. Western blot of senataxin expression in HPV+ parental and resistant cells, following 5□M cisplatin treatment for 24 hours compared to vehicle only control. B. Quantification of three repeats of A (n=3, mean +/- sem). C. Senataxin mRNA expression in untreated HPV+ parental and resistant cells (n=3, mean +/- sem). D. Western blot of senataxin expression in HPV- parental and resistant cells, following 5□M cisplatin treatment for 24 hours compared to vehicle only control. E. Quantification of three repeats of D (n=3, mean +/- sem). F. Senataxin mRNA expression in untreated HPV- parental and resistant cells (n=3, mean +/- sem). G. Cell viability in HPV+ resistant cells in response to cisplatin in presence of senataxin knockdown compared to control (scrambled) siRNA (n=3, mean +/- sem). IC50 for control siRNA: 104.6□M, IC50 for senataxin siRNA: 63.41□M. H. Confirmation of knockdown in HPV+ resistant cells with western blot. I. Confirmation of knockdown in HPV+ resistant cells using qPCR (n=3, mean +/- sem). J. Cell viability in HPV- resistant cells in response to cisplatin in presence of senataxin knockdown compared to control (scrambled) siRNA (n=3, mean +/- sem). IC50 for control siRNA: 96.93□M, IC50 for senataxin siRNA: 66.37□M. K. Confirmation of knockdown in HPV- resistant cells with western blot. L. Confirmation of knockdown in HPV- resistant cells using qPCR (n=3, mean +/- sem). Statistical analysis carried out in B and E using one way-ANOVA. Statistical analysis carried on C, F, I and L using unpaired t-test (with Welch’s correction in I and L). Statistics carried out in G and J using non-linear regression and extra sum-of-squares F test to compare LogIC50 between datasets. ns=not significant, * = p<0.05, ** = p≤0.01, *** = p≤0.001, **** = p≤0.0001.

### Both HPV+ and HPV- resistant cells are sensitised to cisplatin upon senataxin depletion

Dysregulation of senataxin expression has not previously been investigated in the context of cisplatin resistance, which led us to investigate if reduction of senataxin protein levels could modulate the response to cisplatin in resistant cells. Senataxin siRNA led to a successful reduction in protein and mRNA levels in HPV+ resistant cells (Figure 4H-I) and HPV- resistant cells (Figure 4K-L). In the HPV+ resistant cells, depletion of senataxin led to a marked reduction in cell viability in response to cisplatin (Figure 4G). Intriguingly, depletion of senataxin in the HPV- resistant cells also led to a reduction in cell viability following treatment with cisplatin, however the effect was not as marked as in the HPV+ resistant cells (Figure 4J).

### Depletion of senataxin leads to elevated DNA double strand breaks and R-loops following cisplatin treatment

After identifying that depletion of senataxin affected sensitivity to cisplatin in both HPV+ and HPV- resistant cells, we next went onto explore the effect of senataxin depletion on DNA damage. In both HPV+ and HPV- cells, ψH2AX immunofluorescence, revealed an increase in DNA damage following treatment with cisplatin, however the increase in the senataxin depleted cells was significantly greater (Figure 5A-C). Successful knockdown was confirmed using senataxin immunofluorescence (Supplementary Figure 3A-D). There was no increase in DNA damage in the presence of senataxin knockdown alone. These findings were confirmed with an alkaline comet assay under the same conditions in both HPV+ and HPV- resistant cells (Supplementary Figure 2A-B). Following these observations, we next examined whether the reduced cell viability and associated increase in DNA damage noted in the presence of senataxin knockdown was mediated through alterations in R-loop levels. As seen in Figure 5E-H, after cisplatin treatment in senataxin depleted HPV+ resistant cells, there was an increase in R-loop occupancy at certain genomic loci, such as Actin 5’ pause.

**Figure 5:**
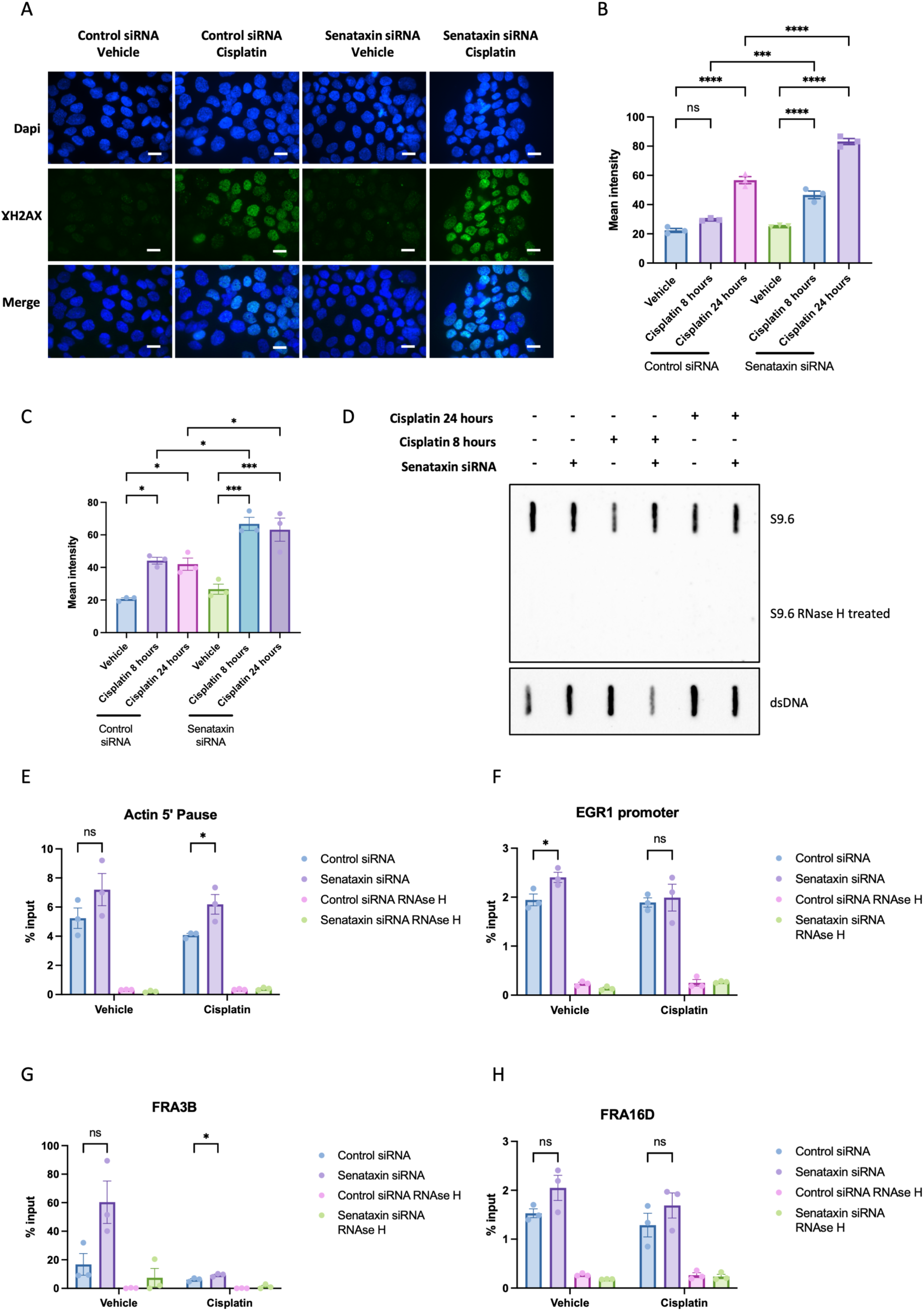
Knockdown of senataxin results in increased DNA damage and R-loops following cisplatin treatment. A. Representative images of Gamma H2AX immunofluorescence after 24 hours of 25□M cisplatin treatment compared to vehicle only control in HPV+ resistant cells. Scale bar 10□M. B. Quantification of mean nuclear intensity following senataxin knockdown and 25□M cisplatin treatment in HPV+ resistant cells for 8 or 24 hours as indicated (n=3, mean +/- sem). C. Quantification of mean nuclear intensity following senataxin knockdown and 25□M cisplatin treatment in HPV- resistant cells for 8 or 24 hours as indicated (n=3, mean +/- sem). D. S9.6 slot blot to investigate global changes in R-loops following senataxin knockdown and 25□M cisplatin treatment for 8 and 24 hours. E. DRIP-qPCR following senataxin knockdown and vehicle or cisplatin treatment at Actin 5’ Pause locus (n=3, mean +/- sem). F. DRIP-qPCR following senataxin knockdown and vehicle or cisplatin treatment at EGR1 promoter locus (n=3, mean +/- sem). G. DRIP-qPCR following senataxin knockdown and vehicle or cisplatin treatment at FRA3B locus (n=3, mean +/- sem). H. DRIP-qPCR following senataxin knockdown and vehicle or cisplatin treatment at FRA16D locus (n=3, mean +/- sem). Statistical analysis carried out on B and C with one-way ANOVA and multiple comparisons carried out using post-hoc Tukey’s test. Statistics carried out on E-H using unpaired t-tests. ns=not significant, * = p<0.05, ** = p≤0.01, *** = p≤0.001, **** = p≤0.0001.

This correlated with a global increase in R-loops under the same conditions (Figure 5D).

### USP11 re-sensitises HPV- resistant cells to cisplatin

Recently, USP11 has been identified as a novel R-loop regulator, acting through de- ubiquitination of senataxin, thus reducing its degradation ^55^. Upon development of resistance in HPV+ cells there was a significant reduction in USP11 protein expression (Figure 6A-B), however there was no significant difference in mRNA level (Figure 6C). In keeping with the lack of change in senataxin expression, there was no significant difference in USP11 expression in HPV- resistant cells (Figure 6D-F). The observed down-regulation of USP11 in HPV+ resistant cells may be a cellular response to the increased expression of senataxin. Reduction in USP11 protein expression has been shown to effect R-loop homeostasis through its effect on senataxin ubiquitination, therefore we next investigated whether targeting USP11 could re-sensitise resistant cells to cisplatin. Depletion of USP11 in HPV+ resistant cells did not affect cell viability in response to cisplatin treatment (Figure 6G). However, in the HPV- resistant cells, depletion of USP11 led to reduced cell viability following cisplatin treatment (Figure 6I). In addition, in HPV- resistant cells, knockdown of USP11 led to increased DNA damage, as measured by ψH2AX immunofluorescence (Figure 6K-L) and alkaline comet assay respectively (Supplementary Figure 2C). Furthermore, in HPV- resistant cells, depletion of USP11 with siRNA led to an associated reduction in senataxin protein following cisplatin treatment (Supplementary Figure 3G-I).

**Figure 6:**
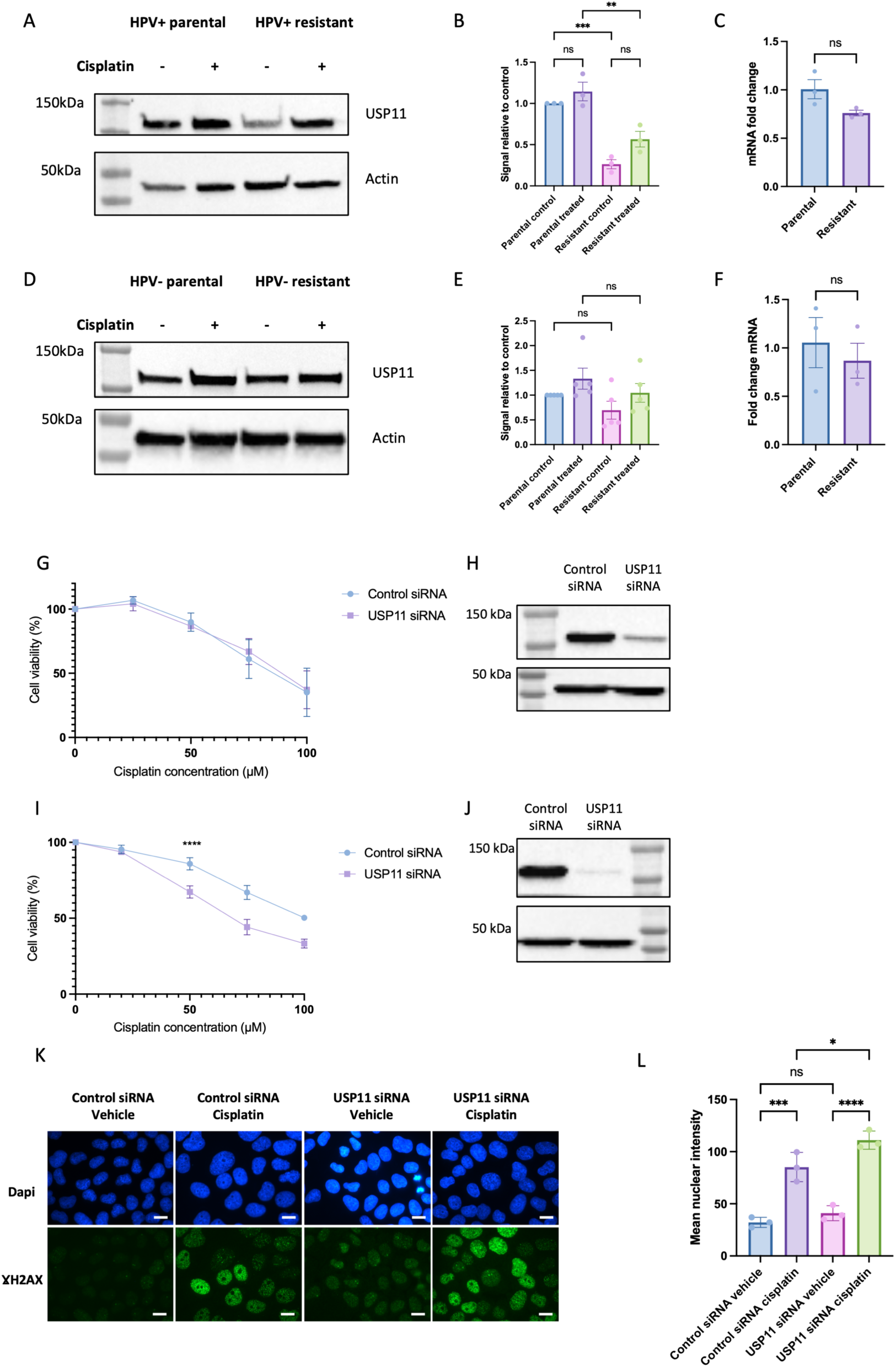
USP11 re-sensitises HPV- resistant cells to cisplatin. A. Western blot of USP11 expression in HPV+ parental and resistant cells, following 5□M cisplatin treatment for 24 hours compared to vehicle only control. B. Quantification of three repeats of A (n=3, mean +/- sem). C. USP11 mRNA expression in untreated HPV+ parental and resistant cells (n=3, mean +/- sem). D. Western blot of USP11 expression in HPV- parental and resistant cells, following 5□M cisplatin treatment for 24 hours compared to vehicle only control. E. Quantification of three repeats of D (n=5, mean +/- sem). F. USP11 mRNA expression in untreated HPV- parental and resistant cells (n=3, mean +/- sem). G. Cell viability in HPV+ resistant cells in response to cisplatin in presence of USP11 knockdown compared to control (scrambled) siRNA (n=3, mean +/- sem). IC50 for control siRNA: 84.97□M, IC50 for USP11 siRNA: 88.09□M, p=0.7476. H. Confirmation of USP11 knockdown at protein level with western blot. I. Cell viability in HPV- resistant cells in response to cisplatin in presence of USP11 knockdown compared to control (scrambled) siRNA (n=3, mean +/- sem). IC50 for control siRNA: 100.6□M, IC50 for USP11 siRNA: 69.59□M. J. Confirmation of USP11 knockdown at protein level with western blot. K. Representative images of Gamma H2AX immunofluorescence after USP11 knockdown in HPV- resistant cells compared to control (scrambled) siRNA and 24 hours of cisplatin treatment compared to vehicle only control. Scale bar 10□M. L. Quantification of mean nuclear intensity in K (n=3, mean +/- sem). Statistical analysis carried out in B, E and L using one way-ANOVA with post-hoc Tukey’s test. Statistical analysis carried on C and F using unpaired t-test. Statistics carried out in G and I using non-linear regression and extra sum-of-squares F test to compare LogIC50 between datasets. ns=not significant, * = p<0.05, ** = p≤0.01, *** = p≤0.001, **** = p≤0.0001.

### In OPSCC tumours which have metastasised to bone, there is evidence of increased R- loops levels by S9.6 immunohistochemistry

To explore whether these findings may be replicated *in vivo*, we sought to investigate the R-loop burden using S9.6 immunohistochemistry in a cohort of 17 HPV+ and HPV- tumours which had responded poorly to treatment. The HPV status of these tumours was confirmed using HPV DNA in-situ hybridisation (ISH) and the specificity of the signal was confirmed with RNase H treatment (Supplementary Figure 4). Following quantification of the S9.6 immunohistochemistry for each of tumours, it was identified that HPV+ tumours have significantly higher S9.6 expression compared to HPV- tumours, however there was no correlation of S9.6 expression with other clinical characteristics including gender, tumour stage, nodal stage, smoking status or alcohol history (Figure 7A-F). Within the cohort, there were samples from different sites, including primary tumour, soft tissue metastases and bone metastases. When the S9.6 expression was compared between these groups, a higher mean S9.6 H-Score was observed in the bone metastases when compared to the primary tumours and soft tissue metastases (Figure 7G-K).

**Figure 7:**
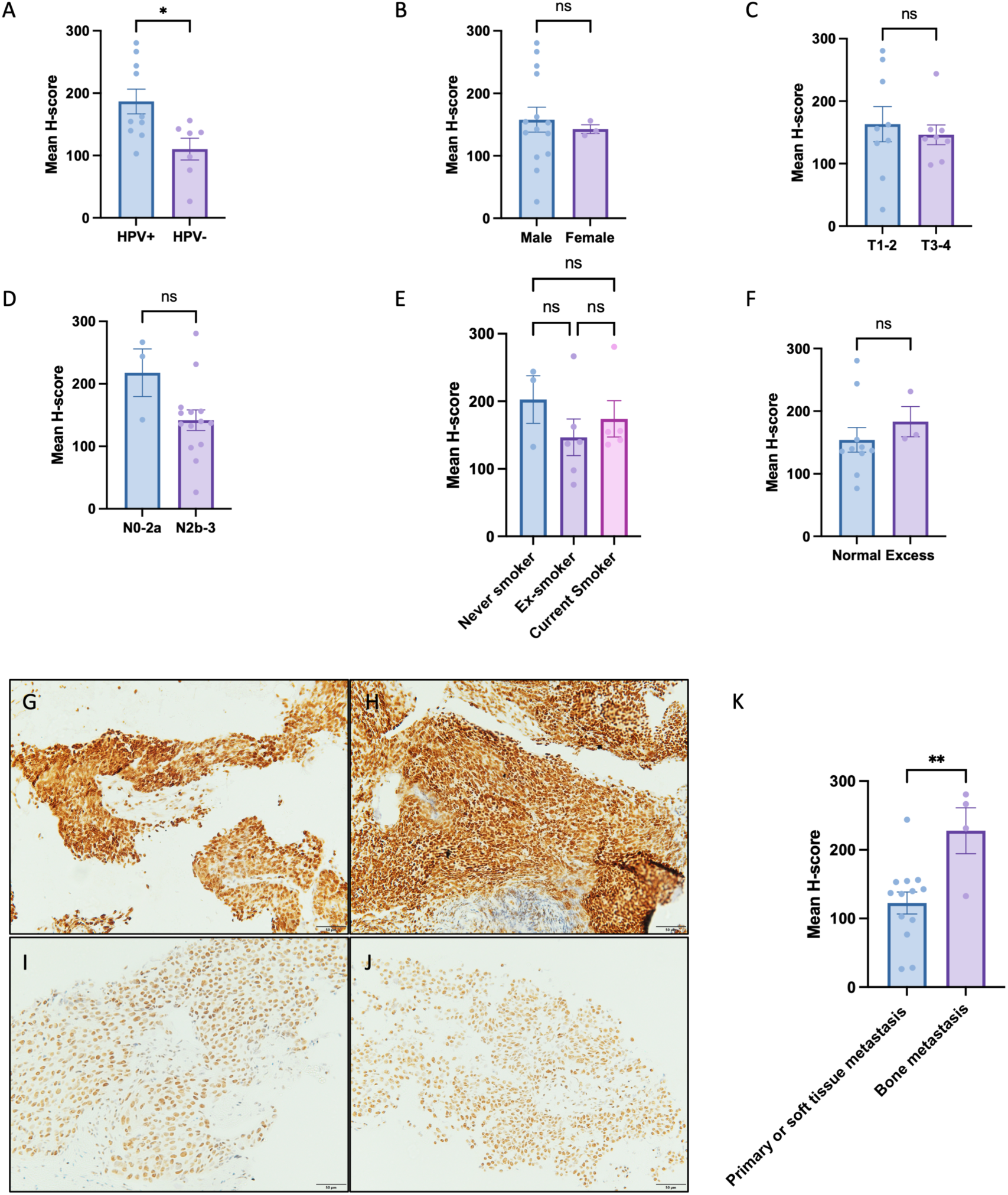
Immunohistochemistry demonstrates higher S9.6 expression in HPV+ tumours and bone metastases. A. Mean H-score by HPV status (n=17, mean +/- sem). B. Mean H-score by Gender (n=17, mean +/- sem). C. Mean H-score by Tumour stage (n=17, mean +/- sem). D. Mean H-score by Nodal stage (n=17, mean +/- sem). E. Mean H-score by smoking status (n=14, mean +/- sem). F. Mean H-score by recorded alcohol consumption (n=14, mean +/- sem). 3 cases excluded from E and F due to missing clinical data. G-J. Representative images of primary tumours, soft tissue metastases and bone metastases. G-H. Representative image of S9.6 staining in bone metastases. I. Representative image of S9.6 staining in soft tissue (lymph node) metastasis. J. Representative image of S9.6 staining in primary biopsy (tonsil). K. Quantification of H-score in primary tumours and soft tissue metastasis compared to bone metastasis. Statistical analysis carried out in A-D, F and K using unpaired t-test. Statistics carried out in E using one-way ANOVA with post-hoc Tukey’s test. ns= not significant,* = p<0.05, ** = p≤0.01.

## Discussion

In this paper, we have demonstrated that in HPV+ cells, resistance to cisplatin can be modulated by senataxin in an R-loop mediated manner. Development of a cisplatin resistant HPV+ and HPV- cell line uncovered new insights into the biology of platinum resistance in head and neck cancer. Novel gene changes in gene expression upon development of cisplatin resistance in head and neck cancer were identified, including CNGB1 and KRT6A. CNGB1 has been associated with cisplatin resistance and reduced progression free survival in bladder cancer ^44^, but has not previously been reported to be involved in development of platinum resistance in HPV- head and neck cancer. Upon development of resistance in the HPV+ cells, overexpression of KRT6A was observed, and this previously been reported in a cisplatin resistant variant of a gastric adenocarcinoma cell line ^56^. Interestingly, KRT6A was down-regulated in the HPV- resistant cells, suggesting HPV+ and HPV- cells have differing gene expression profiles upon development of cisplatin resistance.

In both HPV+ and HPV- cells, development of cisplatin resistance was associated with differential expression of genes related to oxidative phosphorylation. As discussed above, cisplatin treatment is known to increase the levels of metabolites associated with generalised oxidative stress ^46–48^. It has been shown that DNA damage induced by cisplatin can be regulated through modulation of oxidative stress and that ROS scavengers (such as N-acetyl cysteine (NAC)) reduces both the levels of ψH2AX in response to cisplatin and associated apoptosis ^46,49^. Furthermore, Yu *et al.,* demonstrated that development of two cisplatin resistance clones resulted in differential gene expression with an enrichment in genes associated with oxidative phosphorylation ^46^.

After observing that oxidative phosphorylation pathways were differentially expressed upon development of resistance and recognising that ROS are known to increase R-loop levels ^50–52^, we next went onto explore if there was differential expression of R-loop regulators upon development of resistance. Interestingly, we observed differential expression of R-loop regulators in both HPV+ and HPV- resistant cells. To the best of our knowledge, this is the first time that differential expression of R-loop regulators has been explored in the context of cisplatin resistance.

To further understand if R-loops may play a role in the development of cisplatin resistance, we explored R-loop burden in these cells, both globally and at specific loci. Using an S9.6 slot blot, we identified that global R-loop levels increased following cisplatin treatment in cisplatin sensitive HPV+ and HPV- cells (Figure 3A-D). This is in agreement with the literature which shows that DNA crosslinking agents, such as mitomycin C ^57^ and formaldehyde ^58^, increase global R-loop levels. It has been hypothesised that double strand breaks cause stalling of the transcription machinery, which allows the nascent mRNA to thread back between the DNA strands ^59^. In keeping with this theory, the pausing of RNA polymerase II has been shown to increase R-loop formation at transcription termination sites ^60^.

Therefore, as cisplatin creates bulky intra- and inter-strand crosslinks, it is plausible to predict that the presence of cisplatin crosslinks causes pausing of the transcription machinery, leading to increased R-loop levels. To the best of our knowledge, there are no papers in the literature that have shown an increase in R-loops following cisplatin treatment in human cells. There is a single study in mice that investigated the role of R-loops during mouse zygote genome activation ^61^ and identified changes in R-loop dynamics throughout this process, which were linked to transcription ^61^. Interestingly, they treated mouse embryos with cisplatin and doxorubicin and noted an increase in ψH2AX levels by

immunofluorescence, but no significant difference in R-loops levels as measured by S9.6 immunofluorescence following cisplatin treatment for 8 hours ^61^. This contrasts with the findings here; however, it is likely that findings differ due to biological differences between mouse embryos and human cancer cells, alongside the different time points and methodology used in this study. A broadly similar increase in global R-loops was observed in HPV+ resistant cells following treatment with cisplatin, however HPV- resistant cells did not show an increase in R-loops following treatment. It is plausible to assume that these cells are somehow managing to overcome the effects of cisplatin treatment, either by not forming as many cisplatin crosslinks initially or by being able to remove them more effectively.

When comparing differences between parental and resistant cells in untreated conditions, we identified a global increase in R-loops in untreated conditions upon development of cisplatin resistance in both HPV- and HPV+ cells. Increased R-loop levels have been shown to be associated with other aggressive tumours such as Embryonal Tumour with Multilayered Rosettes (ETMR) ^30^ and Ewing sarcoma ^31^. There are several potential explanations as to why R-loops may increase upon development of cisplatin resistance. The first potential explanation is the relationship between R-loops and genomic instability. For example, ETMR has been shown to be genomically structurally unstable with overlap of R-loop signal with regions which have copy number alterations, suggesting R-loops may be a precursor to chromosomal breaks ^30^. Therefore, one possible explanation is that upon development of cisplatin resistance, the cells become more genomically unstable, and this may be driven by increased R-loops. In support of this theory, two studies analysed cisplatin resistant ovarian cancer cell lines and their sensitive parental counterparts and identified copy number alterations with gains and loss identified across all chromosomes ^62,63^, alongside DNA translocations ^63^. Genome wide experiments, such as whole genome sequencing (WGS) and DRIP-Seq could be used to investigate these findings further. It is also conceivable that the resistant cells may be utilising R-loops to modify the chromatin landscape and alter gene expression in a favourable manner which aids resistance. R-loops have been shown to be associated with both transcriptional silencing ^64^ and activation ^65,66^. This could be explored further using genome wide R-loop mapping techniques to investigate R-loop occupancy at the promoters of genes known to be differentially expressed and involved in cisplatin resistance.

Alongside an increase in R-loops globally and at specific loci, HPV+ cells also upregulated senataxin upon development of resistance and depletion of senataxin led to increased sensitivity to cisplatin, increased DNA damage, and increased R-loops both globally and at specific loci. This is in agreement with a genome wide CRISPR screen which showed that loss of senataxin led to sensitised RPE1 cells to cisplatin ^67^. Senataxin has also been shown to be important in resolving R-loops at DNA DSBs ^68^ and depletion of senataxin reduced cell survival following induced DSBs ^68^. Furthermore, there were increased levels of ψH2AX and 53BP1 following senataxin knockdown and induced DSBs, whereas Rad51 levels were reduced ^68^. Therefore, it could be hypothesised that in the setting of cisplatin resistance, increased levels of senataxin would allow for increased resolving of R-loops formed at DNA DSBs, allowing Rad51 recruitment and improved cell viability.

In addition, we demonstrated that senataxin can be targeted through USP11, which is consistent with what has been previously shown ^55^. However, reduced cell viability in the presence of USP11 depletion was only observed in HPV- resistant cells which do not down-regulate USP11. Finally, using S9.6 immunohistochemistry we showed that bone metastases have higher R-loop levels when compared to primary tumours or soft tissue metastases.

We acknowledge a number of limitations, including the cell line model of cisplatin resistance used. A cell line model was chosen as it provided the ability to create cisplatin resistant cell lines and explore the differences between the cisplatin sensitive and resistant cells. It has been shown that lymph node metastasis from OPSCC are heterogeneous in nature ^69,70^, therefore, evaluation of single clones may not be representative of the complexity of cisplatin resistance *in vivo*. However, the model used in this paper allows for manipulation of R-loops that would be impossible in clinical samples. Future work could include evaluating R-loops in tissue samples alongside other markers known to be important in tumour progression (such as immune response and DNA repair), to determine if R-loops are found uniformly throughout a tumour and whether there is any association with other markers which may contribute to treatment resistance. The S9.6 immunohistochemistry was carried out on a relatively small and retrospective cohort. In addition, the findings cannot be directly compared with the cell culture data, as for each of the bone metastases there was only these samples to analyse, and the primary tumour was not available for analysis. However, these findings warrant further investigation to determine if the high level of R-loops in bone metastases may be a potential therapeutic target.

In summary, in this study we have shown that senataxin modulates cisplatin resistance through an R-loop mediated mechanism in HPV+ OPSCC, with potential therapeutic benefits for patients’ who develop cisplatin resistance.

## Acknowledgements

The authors would like to thank Dr Bernadette Foran for her help in collating the clinical cohort and Hayley Stanhope for her technical assistance.

## Funding sources

This study was kindly funded by a Wellcome Trust 4ward North Academy Clinical PhD Fellowship (R120782), a CRUK/Pathological Society Predoctoral Research Bursary (C66701/A27282) and a Pathological Society Equipment Grant (EG20201242). Sherif El-Khamisy is supported by a Wellcome Trust Investigator Award (103844) and a

Lister Institute of Preventative Medicine Fellowship (137661).

## Author contributions

Conceptualization: HC, KDH, SEK; Methodology, HC, IC, KDH, SEK; Validation: HC; Formal analysis: HC; Investigation: HC; Resources: SEK, KDH; Data curation: HC; Writing - original draft: HC; Writing - review & editing: HC, IC, SEK, KDH; Supervision: SEK, KDH; Project administration: HC, SEK, KDH; Funding acquisition: HC, KDH, IC, SEK.

## Conflict of interest

None of the authors have a conflict of interest to declare

## Supplementary material

**Table 1:**
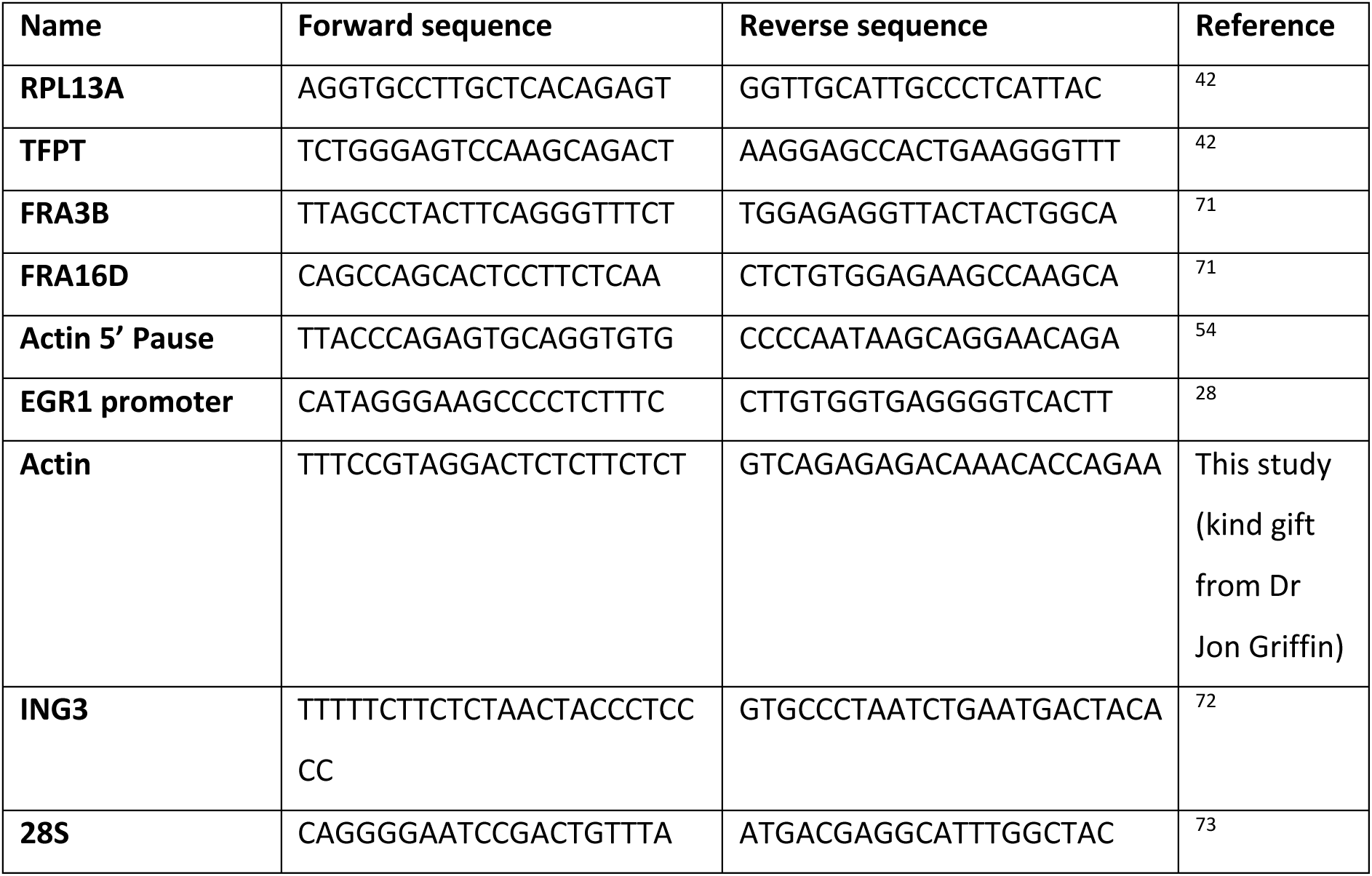
DRIP-qPCR primers used in this study.

**Table 2:**
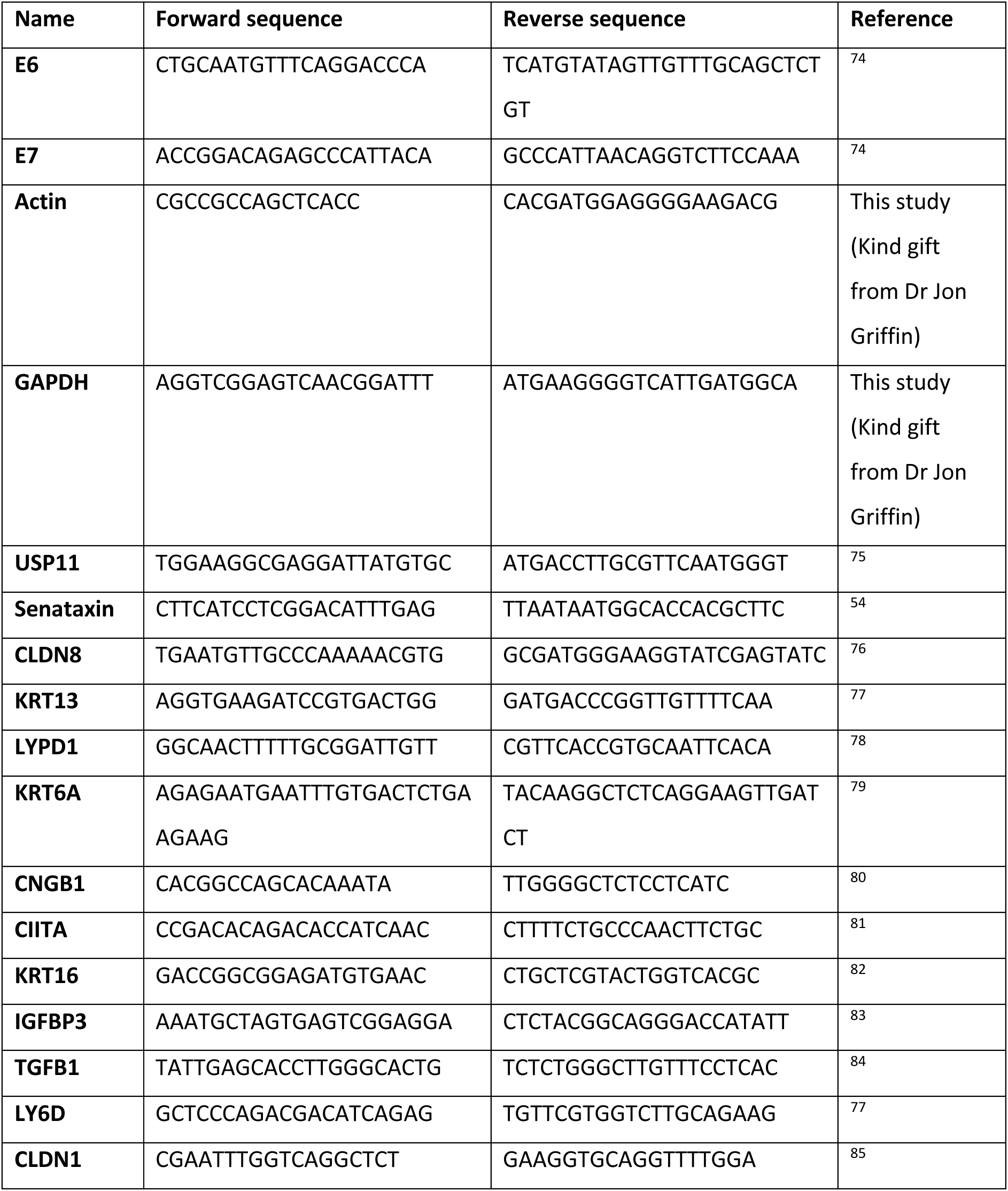
RT-qPCR primers used in this study.

**Table 3:**
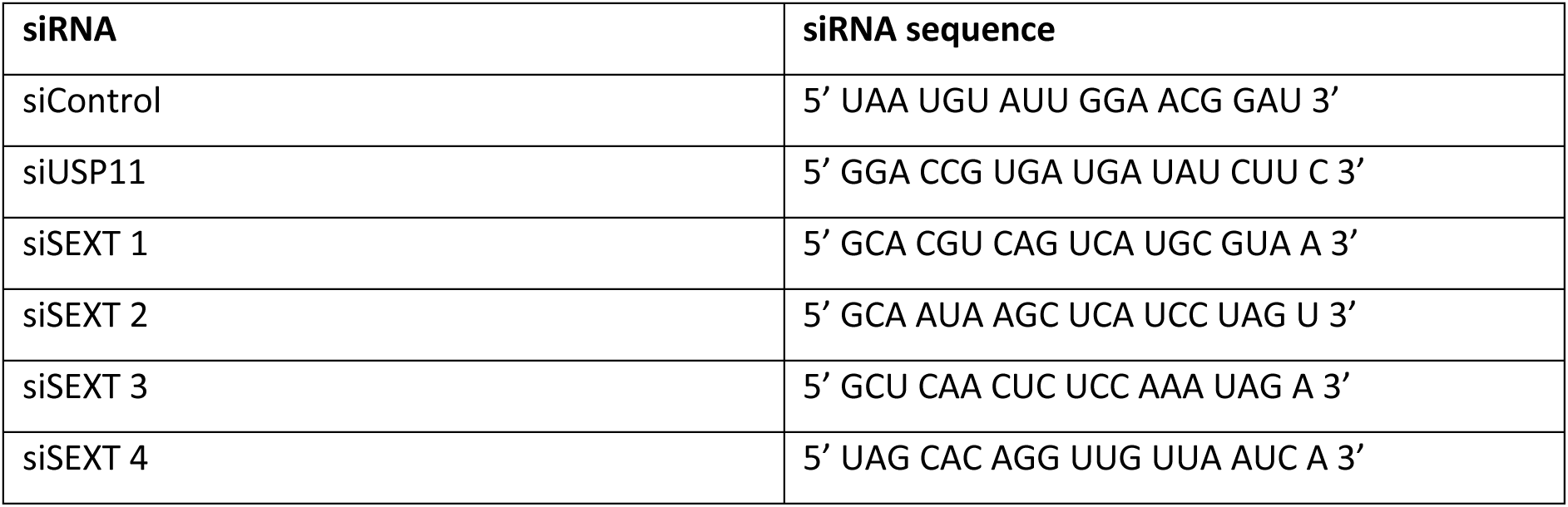
siRNA sequences.

**Supplementary Figure 1:**
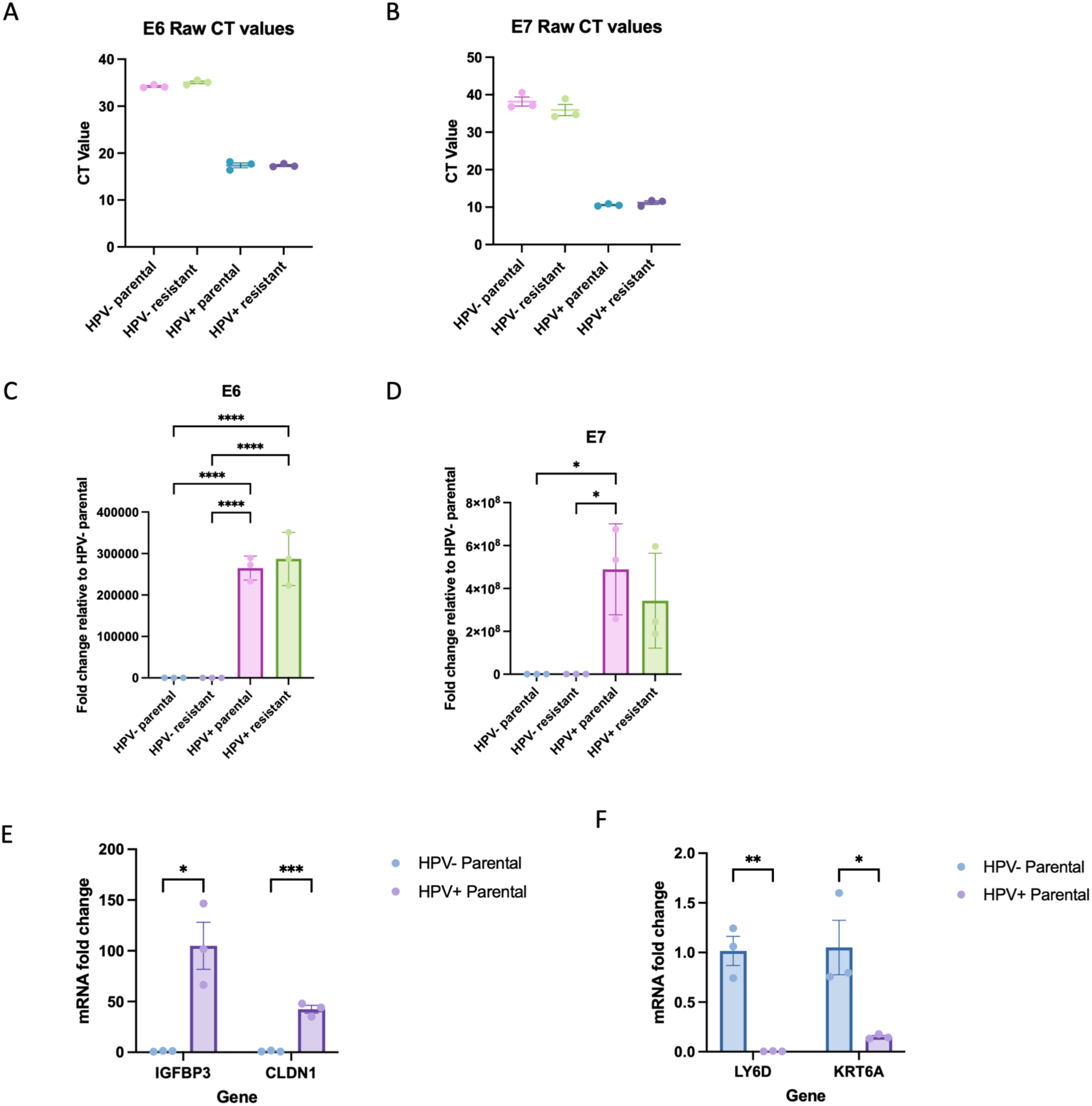
qPCR for E6/E7 viral transcripts and validation of RNA-Sequencing analysis confirms the HPV status of the cell lines. A. E6 Raw CT values. B. E7 Raw CT values. C. Fold change of E6 expression relative to HPV- parental cells (n=3, mean+/- sem). D. Fold change of E7 expression relative to HPV- parental cells (n=3, mean +/- sem). E. qPCR validation of two up-regulated transcripts in the HPV+ parental cells when compared to the HPV- parental cells. F. qPCR validation of two up-regulated transcripts in the HPV+ parental cells when compared to the HPV- parental cells. Statistical analysis carried out on C and D using one-way ANOVA with Tukey’s post-hoc test. Statistical analysis carried out on E and F using multiple unpaired t-tests. ns=not significant, * = p<0.05, ** = p≤0.01, *** = p≤0.001, **** = p≤0.0001.

**Supplementary Figure 2:**
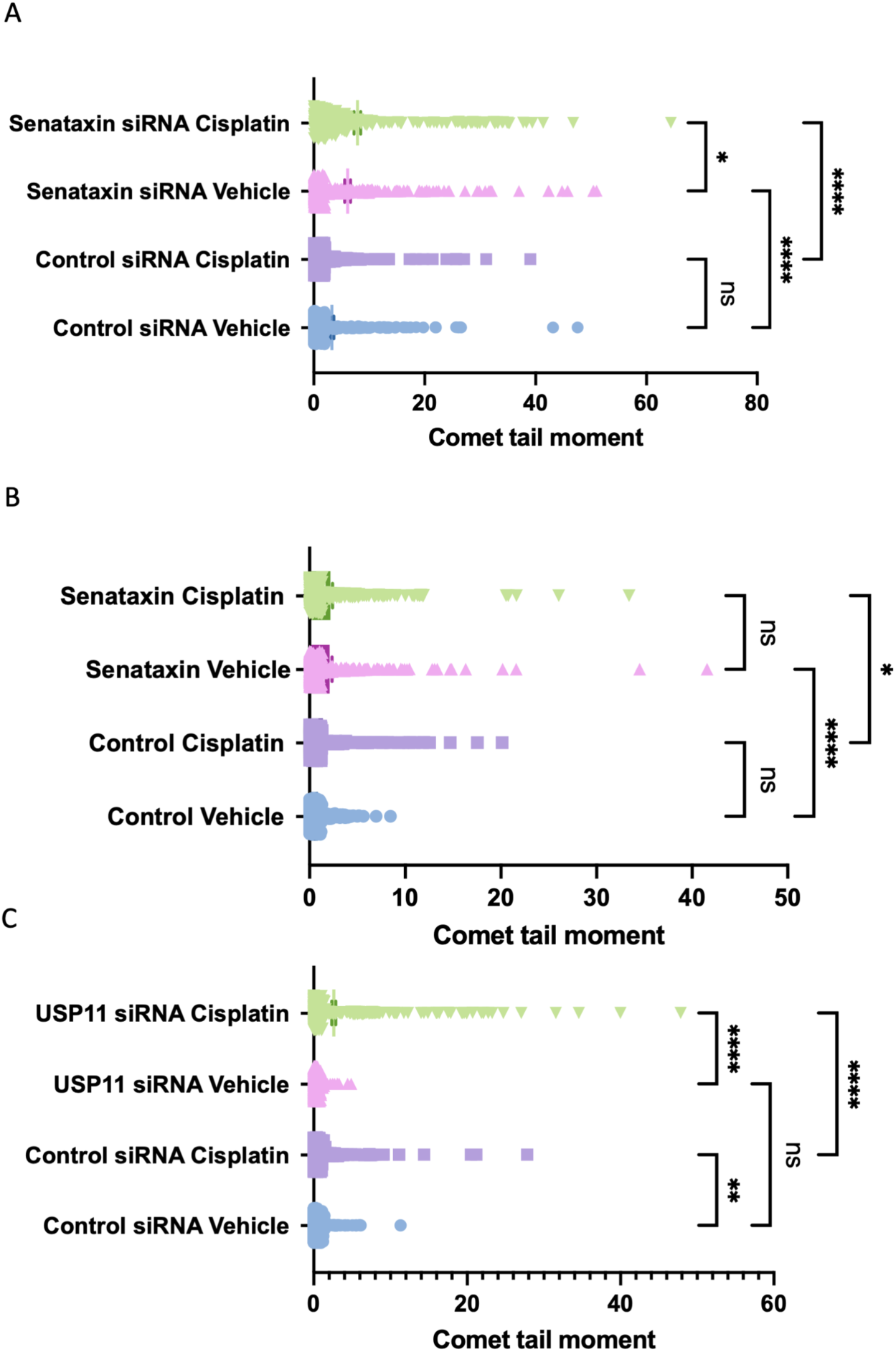
Alkaline single gel electrophoresis (comet) assays confirm increased DNA damage following cisplatin treatment in the presence of senataxin and USP11 depletion. A. Comet assay following senataxin depletion and 24 hours of 25□M cisplatin treatment in HPV- resistant cells. B. Comet assay following senataxin depletion and 24 hours of 25□M cisplatin treatment in HPV+ resistant cells. C. Comet assay following USP11 depletion and 24 hours of 50□M cisplatin treatment in HPV- resistant cells. For all figures, mean +/- sem is plotted alongside all data points from 3 biological replicates, with at least 100 cells counted per biological replicate. Statistical analysis carried out with one-way ANOVA and multiple comparisons carried out using post-hoc Tukey’s test. ns=not significant, * = p<0.05, ** = p≤0.01, *** = p≤0.001, **** = p≤0.0001.

**Supplementary Figure 3:**
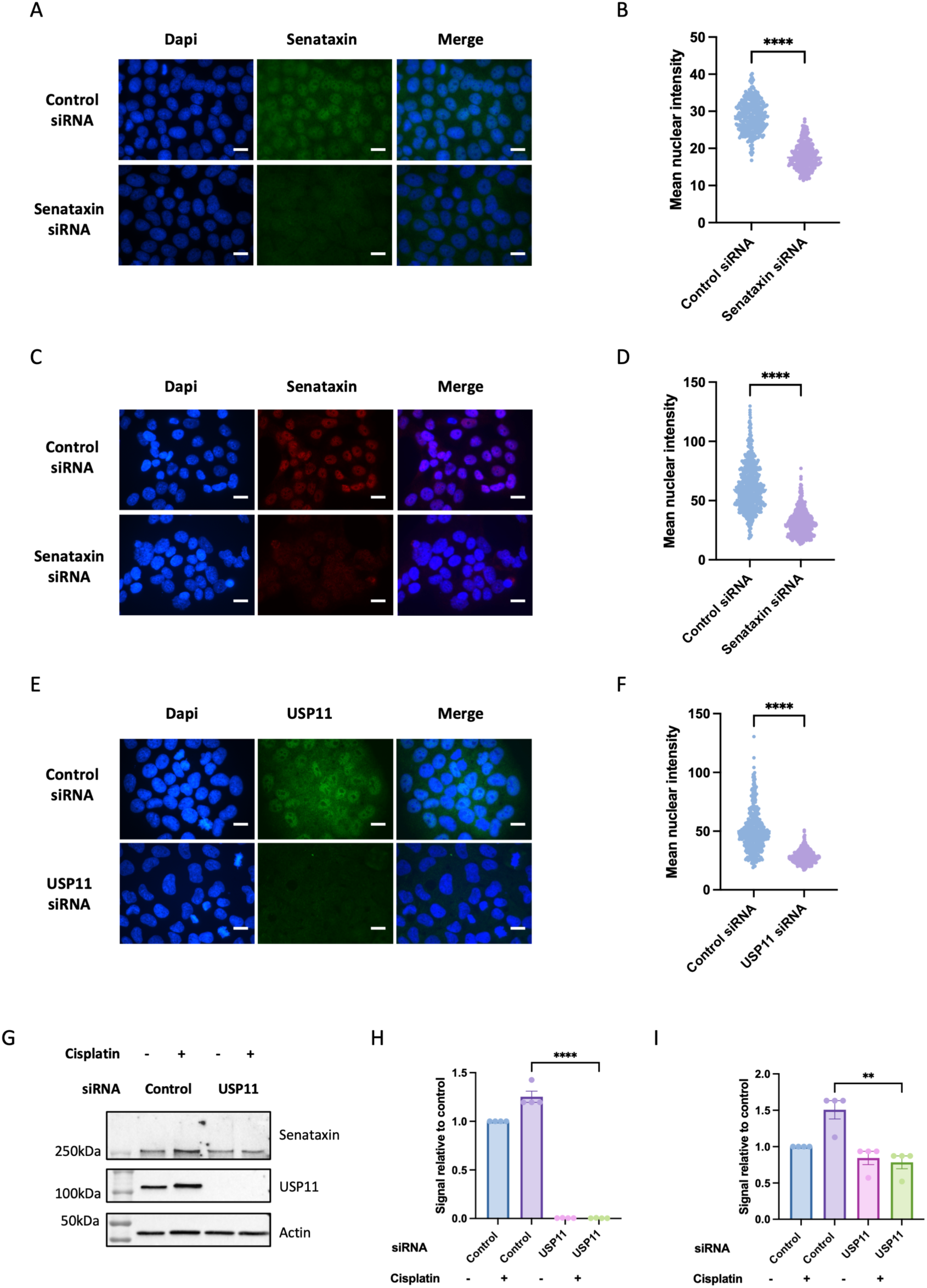
Immunofluorescence to confirm Senataxin and USP11 knockdown and effect of USP11 depletion on senataxin protein expression. A. Representative images of senataxin immunofluorescence in control (scrambled) and senataxin siRNA treated HPV- resistant cells. B. Quantification of senataxin nuclear intensity in control and senataxin siRNA groups from Figure A (all data points from two biological replicates plotted, at least 100 cells quantified per biological replicate). C. Representative images of senataxin immunofluorescence in control (scrambled) and senataxin siRNA treated HPV+ resistant cells. D. Quantification of senataxin nuclear intensity in control and senataxin siRNA groups from Figure C (all data points from three biological replicates plotted, at least 100 cells quantified per biological replicate). E. Representative images of USP11 immunofluorescence in control (scrambled) and USP11 siRNA treated HPV- resistant cells. F. Quantification of USP11 nuclear intensity in control and USP11 siRNA groups from Figure E (all data points from three biological replicates plotted, at least 100 cells quantified per biological replicate). G. Representative western blot of HPV- resistant cells treated with USP11 siRNA or control (Scrambled siRNA) and NaCl vehicle or 50□M of cisplatin for 24 hours. H. Quantification of USP11 expression from G (n=4, mean +/- sem). I. Quantification of senataxin expression from G (n=4, mean +/- sem). Statistical analysis carried out in B, D, F, H and I using unpaired t-test, ** = p≤0.01, **** = p≤0.0001. Scale bar 10□M.

**Supplementary Figure 4:**
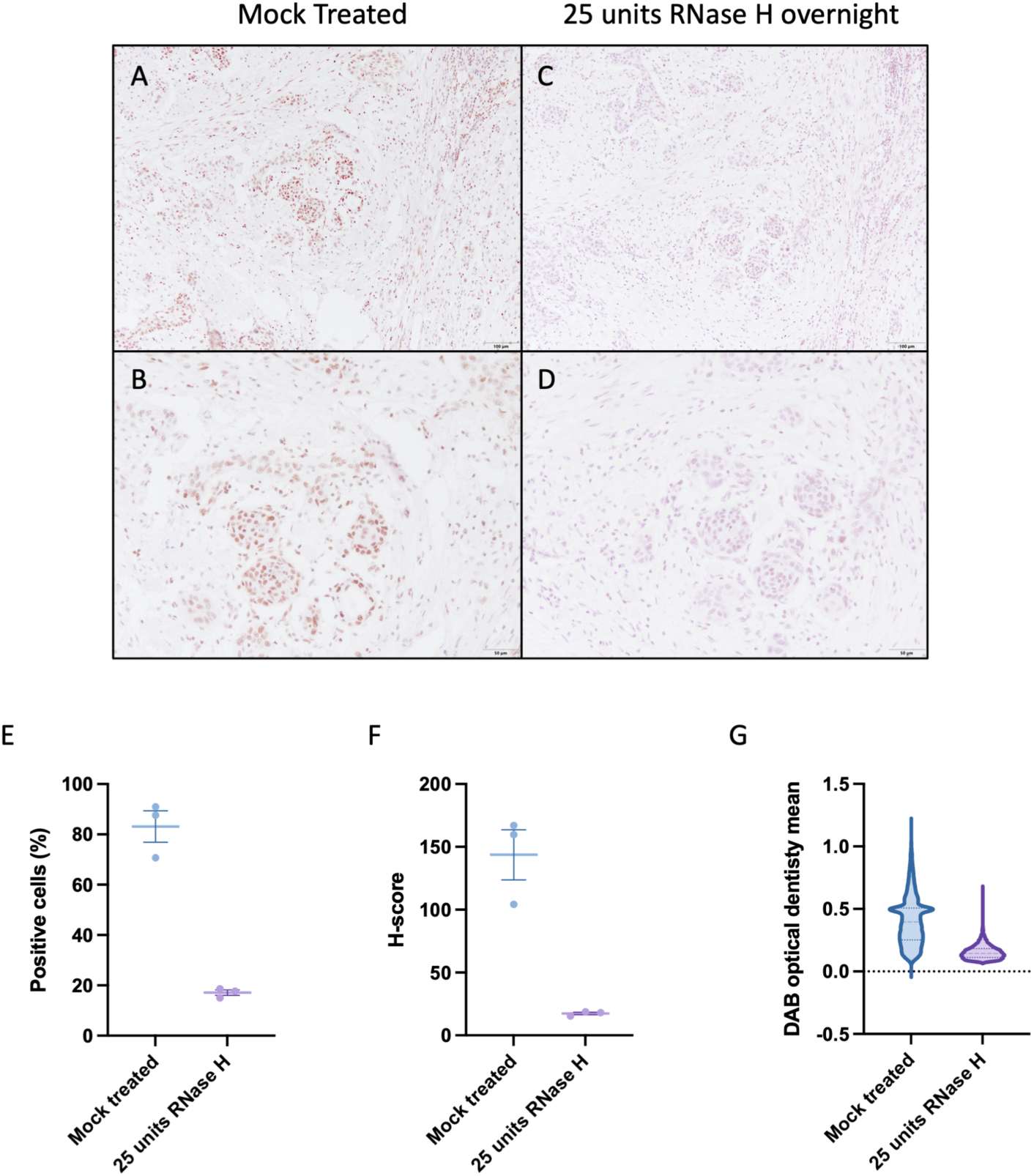
RNase H treatment to confirm specificity of S9.6 immunohistochemistry. A-B. Slides were stained with S9.6 antibody at 1:1000 dilution with mock RNase H treatment (buffer only) overnight. C-D. Serial sections of tissue as A-B stained with S9.6 antibody at 1:1000 dilution with 25 units of RNase H treatment (NEB) overnight prior to blocking. E. Percentage of positive cells from three areas of mock treated slide and slide pre-treated with 25 units RNase H. F. H-Score from three areas of mock treated slide and slide pre-treated with 25 units RNase H. G. DAB optical density mean from three areas of mock treated slide and slide pre-treated with 25 units RNase H.

## Notes

### Competing Interest Statement

The authors have declared no competing interest.

